# Altered structure and dynamics of pathogenic cytochrome c variants correlate with increased apoptotic activity

**DOI:** 10.1101/2020.06.14.151373

**Authors:** Matthias Fellner, Rinky Parakra, Kirstin O. McDonald, Itamar Kass, Guy N.L. Jameson, Sigurd M. Wilbanks, Elizabeth C. Ledgerwood

**Affiliations:** Department of Biochemistry, School of Biomedical Sciences, University of Otago, Dunedin, New Zealand; Department of Biochemistry and Molecular Biology and Victorian Life Sciences Computation Initiative Life Sciences Computation Centre, Monash University, Clayton, VIC 3800, Australia; School of Chemistry, Bio21 Molecular Science and Biotechnology Institute, The University of Melbourne, Parkville, Victoria 3010, Australia

**Keywords:** cytochrome *c*, apoptosis, apoptosome, peroxidase, thrombocytopenia, molecular dynamics, protein structure, alkaline transition

## Abstract

Mutation of cytochrome *c* in humans causes mild autosomal dominant thrombocytopenia. The role of cytochrome *c* in platelet formation, and molecular mechanism underlying the association of cytochrome *c* mutations with thrombocytopenia remains unknown, although a gain-of-function is most likely. Cytochrome *c* contributes to several cellular processes, with exchange between conformational states proposed to regulate changes in function. Here we use experimental and computational approaches to determine whether pathogenic variants share changes in structure and function, and to understand how these changes might occur. We find that three pathogenic variants (G41S, Y48H, A51V) cause an increase in apoptosome activation and peroxidase activity. Molecular dynamics simulations of these variants, and two non-naturally occurring variants (G41A, G41T), indicate that increased apoptosome activation correlates with increased overall flexibility of cytochrome *c*, particularly movement of the Ω loops. This suggests that the binding of cytochrome *c* to apoptotic protease activating factor-1 (Apaf-1) may involve an “induced fit” mechanism which is enhanced in the more conformationally mobile variants. In contrast, peroxidase activity did not significantly correlate with protein dynamics suggesting that the mechanism by which the variants alter peroxidase activity is not related to the conformation dynamics of the hexacoordinate heme Fe state of cytochrome *c* analyzed in the simulations. Recent suggestions that conformational mobility of specific regions of cytochrome *c* underpins changes in reduction potential and the alkaline transition p*K* were not supported. These data highlight that conformational dynamics of cytochrome *c* drives some but not all of its properties and activities.

## Introduction

Cytochrome *c* is a small heme protein with well described roles in the mitochondrial respiratory chain and in the intrinsic apoptosis pathway (1, 2). In the former, cytochrome *c* transfers electrons from complex III to complex IV, and in the latter cytochrome *c* is released from the mitochondrial intermembrane space and binds to cytosolic Apaf-1 to trigger caspase activation. Both of these functions of cytochrome *c* are essential, with knock-out of cytochrome *c* in mice causing embryonic lethality at E8.5 (3), and knock-in of an apoptosis-inactive cytochrome *c* variant causing late embryonic/early perinatal lethality (4). In addition, native cytochrome *c* has low peroxidase activity, which is proposed to be activated during apoptosis catalyzing oxidation of cardiolipin to assist in the release of cytochrome *c* to the cytosol (1, 5). The ability of cytochrome *c* to undertake these various different roles has led to it being referred to as a “moonlighting” protein, with the adoption of a range of conformational states thought to enable multiple activities (1,6,7).

The recent descriptions of naturally occurring mutations in cytochrome *c* raise new questions about the role of cytochrome *c* in biology. Four pathological variants in *CYCS* have been reported^1^; c.132G>A; p.Gly42Ser (8), c.145T>C; pTyr49His (9), c.155C>T; pAla52Val (10) and c.301_303del:p.Lys101del (11). All cause autosomal dominant thrombocytopenia (THC4, OMIM 612004) characterized by a low count of morphologically and functionally normal platelets, with mild or no bleeding symptoms. Detailed studies of subjects with the c.132G>A; p.Gly42Ser variant have shown that the low platelet count is due to an abnormality in platelet formation in the bone marrow (12), however the underlying role of cytochrome *c* in platelet formation and the molecular basis of THC4 remain unknown.

We have previously characterized G41S cytochrome *c* and two additional non-natural variants, G41A and G41T cytochromes *c* (13, 14). G41S cytochrome *c* has an increased ability to activate caspases and increased peroxidase activity compared to WT, whereas G41T is less able to activate caspases than WT but has greater peroxidase activity than G41S. Recently Y48H and A51V cytochromes *c* have been reported to have increased peroxidase activity compared to WT cytochrome *c* (15, 16).

^1^ Note that the amino acid numbering in the formal description of each variant refers to the protein with the initiating methionine as residue 1. Subsequently we refer to the proteins using numbering for the mature protein, i.e. G41S, Y48H and A51V.

We hypothesize that changes common to the pathological variants are likely to be drivers of the THC4 phenotype. Here we used in vitro and in silico analyses to compare the structure, biophysical properties, and activities, of WT, G41S, G41T and G41A cytochromes *c*, with the recently reported Y48H and A51V cytochromes *c*. This has led to a new understanding of how the conformational space available to cytochrome *c* contributes to these properties.

## Results

### Y48H and A51V mutations increase apoptosome activation by cytochrome *c*

Mutation of Gly41 to Ser increases the apoptotic activity of cytochrome *c* in vitro (8). The Y48H mutation increases the apoptotic activity of mouse cytochrome *c* (9). However cytochrome *c*-induced apoptosis is species specific (17) and the impact of the A51V and Y48H mutation on apoptosome activation by human cytochrome *c* has not previously been determined. We now show that the Y48H and A51V mutations both caused an increase in cytochrome *c*-induced apoptosome activation compared to WT (Table 1), using a cell-free human caspase activity assay. As previously reported (14), apoptosome activation by cytochrome *c* was increased with the G41S mutation but decreased with the G41T mutation (Table 1).

**Table 1.** Apoptosome activation and steady state peroxidase activity of cytochrome *c* variants.

|  | apoptosome activation<br>(normalized to WT) | peroxidase activity<br>(M<sup>-1</sup>s<sup>-1</sup>) |
| --- | --- | --- |
| WT | 1 | 6.3 ± 0.3<sup>b</sup> |
| G41S | 1.7 ± 0.4 | 61.3 ± 3.0<sup>b</sup> |
| G41T | 0.10 ± 0.01 | 94.0 ± 4.0<sup>b</sup> |
| G41A | 0.91 ± 0.02<sup>a</sup> | ND |
| Y48H | 1.5 ± 0.4 | ~ 50<sup>c</sup> |
| A51V | 1.4 ± 0.3 | ~ 34<sup>c</sup> |

ND Not determined
<sup>a</sup> Josephs et al (2013) (14)
<sup>b</sup> Parakra et al (2018) (18)
<sup>c</sup> The reactions are not exactly second order in peroxide so these rate constants are approximate

### Y48H and A51V mutations increase the peroxidase activity of cytochrome *c*

Our previous analyses had shown that the G41S, G41T and G41A mutations increase peroxidase activity of cytochrome *c* across a range of pH (13), and we subsequently determined the second order rate constants for WT, G41S and G41T cytochromes *c* at pH 5.4 (18). The peroxidase reaction catalyzed by cytochrome *c* is characterized by an initial lag phase during which the Met80 sulfur atom is oxidized to a sulfoxide, converting the heme Fe to an active pentacoordinate state, followed by a steady state phase (18). Y48H and A51V cytochrome *c* showed similar progress curves with an initial lag phase followed by steady state phase (Figure 1A and B). The dependence of the steady state rate on hydrogen peroxide (H_2_O_2_) was found to be not quite first order (Figure 1C) suggesting a slightly more complicated reaction mechanism, but an approximate second order rate constant was obtained to compare with other variants (Table 1). It can be seen that they are comparable with G41S, and overall we conclude that all pathogenic mutations lead to an approximately tenfold increase in peroxidase activity of cytochrome *c*.

**Figure 1.**
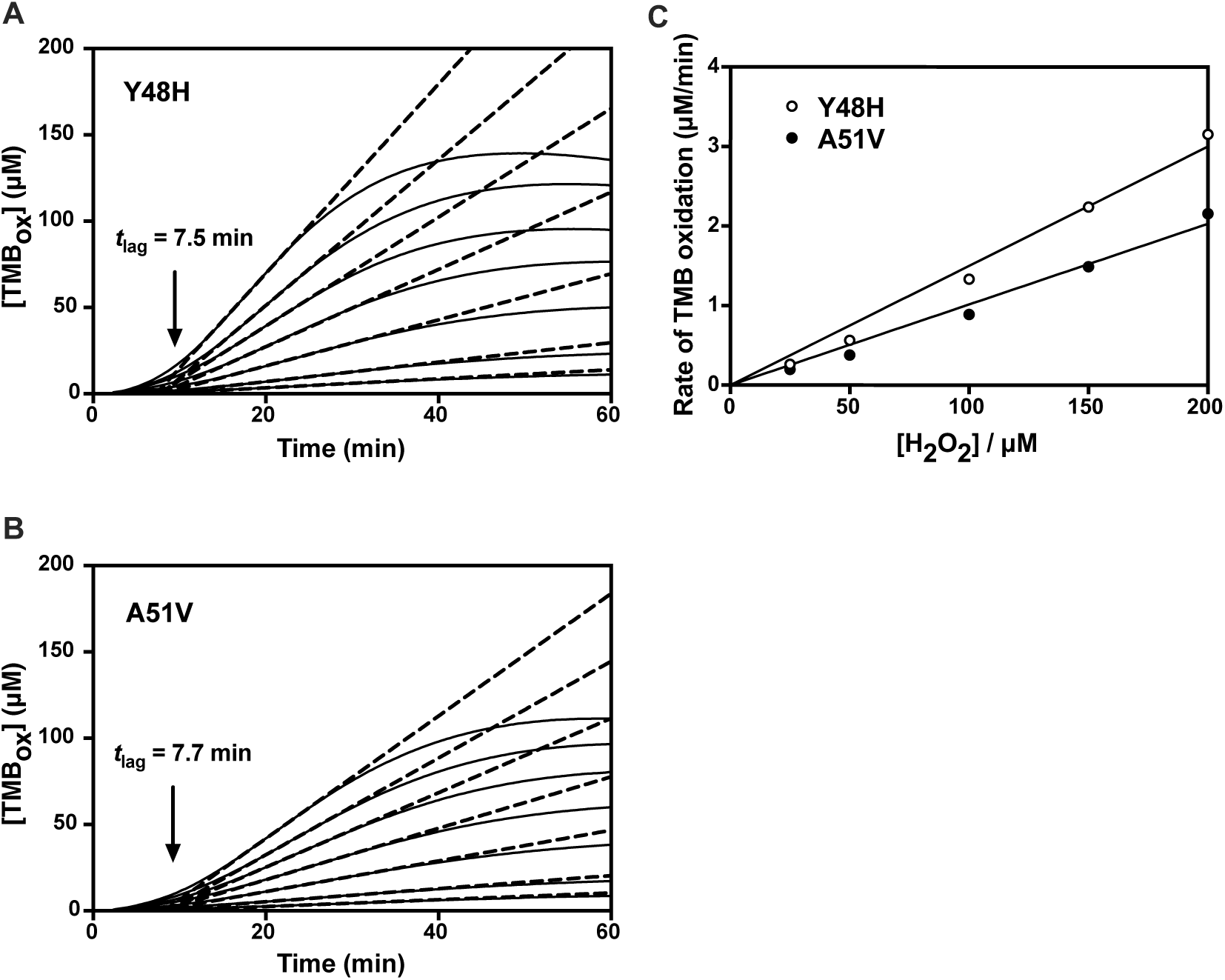
Kinetics of the peroxidase reaction for Y48H and A51V variants of cytochrome *c* with TMB at pH 5.4. (A and B) The plots of time versus [TMB_ox_] show a lag phase followed by a steady state phase that can be fitted to a straight line (dashed). By varying peroxide these lines ([H_2_O_2_] = 25-300 μM) intersect at a single point on the x axis defining the lag time. (C) Plot of the slopes versus [H_2_O_2_] are almost linear, allowing the 2second order rate constant to be approximated (see text). All protein was 5 μM in heme.

### Y48H and A51V mutations decrease the alkaline transition p*K* without altering the redox potential

We and others have previously found that mutation of cytochrome *c* can alter the redox potential, the p*K* of the alkaline transition and the absorbance maximum of the Soret band (14,18,19). We therefore determined these parameters for the new variants (Table S1). The redox potential of all the pathogenic variants is unchanged compared to WT, in keeping with the lack of any phenotype indicative of an alteration in mitochondrial electron transport in subjects with these mutations (20). As previously observed for G41S and G41T cytochromes *c* (18), the maxima of the Soret bands shifted to a slightly lower wavelength in Y48H and A51V cytochromes *c*, indicating a similar change in the Fe spin state (21). The new pathogenic variants show an identical lowering of the p*K* for the alkaline transition (0.9 log units), less than we have previously determined for G41S (1.5 log units) and the non-natural G41T (2.6 log units) or G41A (1.2 log units) variants.

### Mutation of cytochrome *c* causes subtle changes in structure and H-bond networks

To gain insights into the underlying structural basis of variation seen in apoptosome activation, peroxidase activity and biophysical properties occurring due to mutation of cytochrome *c*, we turned to analysis by X-ray crystallography in combination with molecular dynamics (MD) simulations. We determined the crystal structures of the ferrous forms of the Y48H and G41T variants to 1.6 Å and 2.7 Å respectively (SI and Table S2), allowing comparison with previously determined structures of ferric WT cytochrome *c* (19) and ferrous G41S cytochrome *c* (22). Cytochrome *c* is a compact protein, with a core of α helices and β sheets linked by three Ω loops (23) (Figure S1). As previously reported for G41S (22), G41T and Y48H cytochromes *c* have the same fold as WT with a Cα alignment root mean square deviation (RMSD) of 0.3 – 0.4 Å (Figure S2), and a hexacoordinate Fe as expected for the ferrous and ferric forms of the protein.

The G41S, G41T and Y48H mutations are within the 40-to-57 Ω loop and the overall fold of this loop is similar to WT in all crystal structures. However, the G41T and Y48H mutations cause alterations in the hydrogen bond (H-bond) network linking the 40-to-57 Ω loop to the heme (Figures 2A&B, S3) associated with backbone shifts of up to ∼2 Å, similar to our observations in G41S cytochrome *c*. In G41T cytochrome *c*, as previously reported for G41S cytochrome *c* (14), the orientation of the Asn52 sidechain changes. A new H-bond is formed between the Asn52 amide oxygen and the Thr41 hydroxyl (Figure 2A). The Asn52 amide nitrogen H-bonds both directly to heme propionate-7 (hp-7) and also via a water molecule to Thr78. By comparison, in WT cytochrome *c* the amide nitrogen and oxygen participate in one H-bond each, to hp-7 and the water molecule respectively. In Y48H cytochrome *c* (Figure 2B), the Asn52–hp-7 and Asn52–water–Thr78–hp-6 H-bond networks are the same as WT. In the G41S variant, like WT, hp-7 also makes a H-bond to the hydroxyl of Tyr48 which in turn is H-bonded to the amide of Ala43. Although this is no longer possible in Y48H, hp-7 occupies the same position and instead the imidazole ε-N of H48 forms an analogous H-bond to Ala43 while the δ-N H-bonds to hp-6.

**Figure 2.**
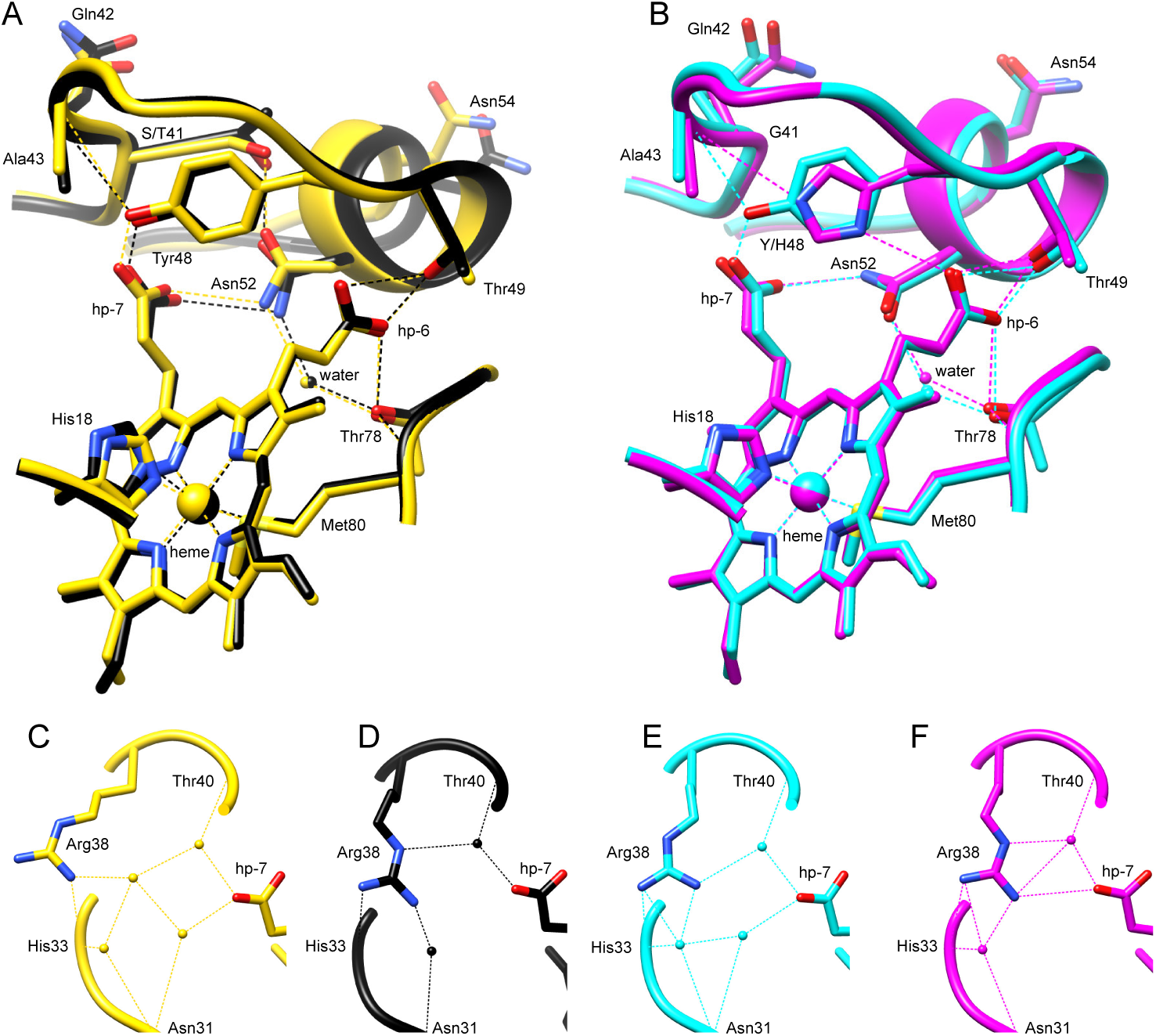
H-bond networks in cytochrome *c* variants. Comparison of the hydrogen bonding network between the human cytochrome *c* 40-57 Ω loop (residues 39-57 shown) and the heme for (A) G41T (black, chain A, 6ECJ) and G41S (yellow, chain B, 3NWV (12)), and (B) WT (cyan, chain A, 3ZCF (13)) and Y48H (magenta, chain A, 5EXQ). Alignment of all structures is in Figure S3. (C-F) H-bond networks linking Arg38 with a heme propionate (hp-7) in each variant (C, G41S; D, G41T; E, WT; F, Y48H).

The impact of the cytochrome *c* mutations on apoptosome activation could be explained if G41S, Y48H and A51V cytochromes *c* have stronger, and G41T cytochrome *c* weaker, affinity for Apaf-1 compared to WT cytochrome *c*. Recent cryo-EM structures of the apoptosome in ground and active states have provided insights into the overall orientation of cytochrome *c* in a binding pocket formed by the WD1 (7 blade β propeller) and WD2 (8 blade β propeller) domains of Apaf-1 (24-26). The resolution of these structures is insufficient to refine mainchain conformation or sidechain orientations of cytochrome *c*. Nonetheless, modelling of the equine cytochrome *c* crystal structure in the cryo-EM electron density identifies five potential polar bonds between the 40-57 Ω loop and the Apaf-1 WD2 domain (Figures 3A and S4).

**Figure 3.**
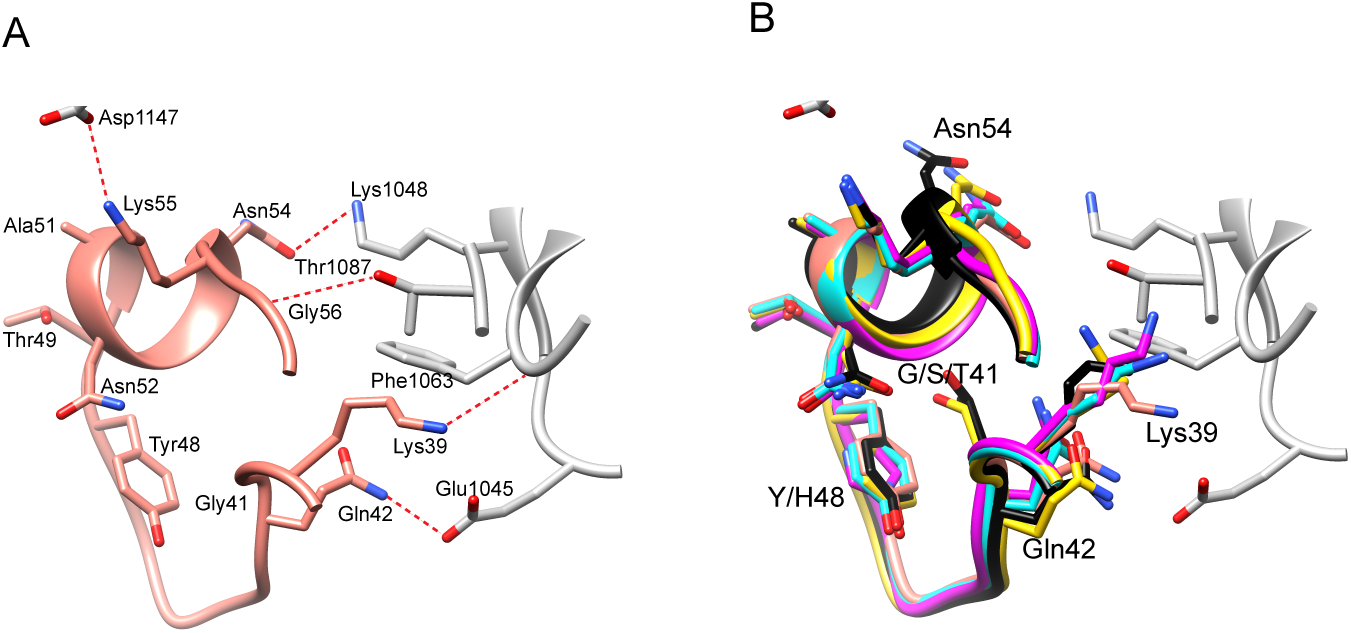
Putative interactions between the 40-57 Ω-loop of cytochrome *c* and the WD2 domain of Apaf-1. (A) Cryo-EM structure of Apaf-1-Caspase-9 holoenzyme (5WVE (16)) showing the docked *Equus caballus* cytochrome *c* (pink, chain B, 4RSZ (17)) interaction with Apaf-1 (grey) in the formed apoptosome. Potential polar bonds between cytochrome *c* and the WD2 domain indicated in red. (B) Superimposed structures of 40-57 Ω-loops of human cytochrome *c* G41T (black, chain A, 6ECJ), G41S (yellow, chain B, 3NWV) WT (cyan, chain A, 3ZCF) and Y48H (magenta, chain A, 5EXQ) onto the *Equus caballus* structure. Apaf-1 in grey.

We superimposed the available structures of human cytochromes *c* on the equine cytochrome *c* structure in the apoptosome complex, to determine whether any differences can be observed which might explain the different interactions of WT and the variants with Apaf-1 (Figure 3B). The crystal structures show Cα movement of up to ∼1.2 Å and 2 Å at the beginning and end of the 40-57 Ω loop respectively (Figure 3B). Variants G41S and G41T show a ∼1.2 Å shift of Gln42 Cα compared to Y48H, with WT occupying an intermediate position. This shift is caused by the differences in the H-bond network at residue 41 and 43 as discussed above (Figure 2). The helical region of the loop shows even greater backbone shifts. There is a difference of 2.0 Å between G41T and WT at residue 54, with these changes potentially being caused by the different side chain orientation of Asn52. The movement of the loop in the variants repositions residues facing outwards towards the WD2 domain (Gln42, Lys39 and Asn54) with the Lys39 sidechain being particularly flexible, modelled differently in all structures. Although there is no consistent pattern of changes in the 40-57 Ω loop, it appears possible that the conformation of this loop is crucial to the interaction between cytochrome *c* and Apaf-1.

The other notable difference between the native and mutant cytochrome *c* structures is the orientation of the Arg38 sidechain (Figure 2C-F). Arg38 lies immediately adjacent to the loop discussed above and is implicated in controlling access of substrate to the heme for the peroxidase reaction (27). We and others have previously pointed out that Arg38 is modelled differently in G41S, with a water molecule in place of the side chain orientation seen in WT (Figure 2C) (13, 19). Additional electron density for the Arg38 sidechain in G41S indicates it has partial occupancy in the WT orientation. In G41T and Y48H cytochromes *c*, which have peroxidase activity similar to or higher than G41S, the Arg38 sidechain position is more similar to WT (Figure 2 D & F) although in some G41T chains, Arg38 could be modelled in an alternative orientation pointing directly towards hp-7 (not shown). Overall, these data suggest that the Arg38 sidechain is likely to be mobile. However, it is not clear whether the mutations alter this mobility, or whether alteration in the orientation of Arg38 explains the increased peroxidase activity of G41S, Y48H and G41T cytochromes *c*.

### Mutation of cytochrome c alters conformational dynamics

The crystal structures show that both the pathogenic and non-natural mutations alter the hydrogen bonding network between the 40-57 Ω loop and the heme. Modelling the structures onto the apoptosome cryo-EM electron density shows that this region makes contacts with Apaf-1. Others have suggested that the mobility of the 40-57 Ω loop influences other Ω loops in cytochrome *c*, and contributes to the impact of the G41S and Y48H mutations on cytochrome *c* functions (7,28,29). Since differences in protein conformational mobility are not captured in the static crystal structures, we next performed MD simulations of WT, G41S, G41T, G41A, A51V and Y48H cytochromes *c*.

There have been numerous MD studies on cytochromes *c* from a range of species, originally focusing on protein folding and unfolding, and more recently on the impact of post-translational modifications and mutations (29-31). Here we analyzed four independent simulations of 200 ns for WT and each variant, using human G41S cytochrome *c* (3NWV) as the starting structure. Apoptosome activation and peroxidase activity are assayed with reduced and oxidized cytochrome *c* respectively. Therefore for each variant two simulations incorporated an oxidized heme (FeIII_1 and FeIII_2) while two incorporated a reduced heme (FeII_1 and FeII_2). In all simulations the protein remained folded (Table S3), and under the parameters used the ligation between Met80 and the heme Fe stays intact (Fe-(S)Met80 bond 2.496 ± 0.002 Å over all simulations), consistent with spectroscopic investigations (14). The 800 ns total simulation analyzed for WT and each variant is the longest yet reported simulation for human cytochrome *c*.

Cytochrome *c* is a relatively small and compact protein so, with the usual caveats (32), root mean square deviation (RMSD) provides a useful initial measure of the effect of mutations on the mobility of the main chain. The RMSD between states obtained during MD and the 3NWV starting structure were plotted and used to construct histograms of the frequency of RMSD values in 0.1 Å increments. Interestingly, this analysis suggested that the physiological variants G41S, Y48H and A51V sample states not well populated by WT, G41A and G41T (Figure S5). These differences in distributions are significant in a Kolmogorov-Smirnov test, although this must be interpreted in the context of the under-sampling inherent in MD simulations of this length (33).

To determine the origin of the RMSD variation and understand differences in mainchain mobility between variants, root mean square fluctuation (RMSF) for Cα of each residue was considered (Figure S6). The main chain Cα for residues in the α-helix and β-sheet regions show little movement (RMSF < 1 Å), further confirming the overall stability of the structures during the simulations. The pattern of main chain movement is similar in WT and variants. Residues with highest Cα RMSF (apart from the extreme N-and C-termini) are within the 19-36 and 40-57 Ω loops, with 8/24 and 16/24 simulations respectively having residues with Cα RMSF > 1 Å (Figure S6). This is consistent with previous MD and hydrogen-deuterium exchange data (7,28,30,34-36). There was no correlation between RMSF in the 19-36 and 40-57 Ω loops (r = 0.28, *P* = 0.18 with two-tailed Pearson correlation), suggesting these loops move independently. There is less movement of the 71-85 Ω loop (3/24 simulations with Cα RMSF > 1 Å). This is expected since the loop is anchored by the Met80-Fe ligation, which is not disrupted under the simulation conditions used.

We next determined the basis of variable RMSF in the Ω loops by examining the changes of distance between critical points in the structure during the simulations. The 40-57 Ω loop is anchored by H-bonds between Tyr/His48, Thr49, Asn52 and the heme. This is part of the H-bond network seen to be altered by mutations in this loop in the crystal structures (Figure 2A and B). Intermediate RMSF are observed when Thr49 and Asn52 lose their connection with the heme, as seen in WT cytochrome *c* after ∼100 ns, A51V cytochrome *c* after ∼30 ns and G41S cytochrome *c* after ∼ 30 ns (Figure 4A-C). As a result, residues 49-57 become independent of the heme, with this part of the loop adopting several conformations for the rest of the simulation (extreme examples shown for WT cytochrome *c* at 192 ns and A51V cytochrome *c* at 150 ns)). Even when the residues 47-57 become mobile, Tyr48 can still keep its heme interaction and anchor the rest of the Ω loop (Figure 4A&B). However, when the Tyr48-heme interaction is also lost, for example at ∼50 ns in G41S cytochrome *c* (Figure 4C), then the entire 40-57 Ω loop can adopt multiple different conformations resulting in RMSF > 2.0 Å for this region (Figure S6C). The 19-36 Ω loop also displays residues with RMSF greater than 2.0 Å in some simulations. The main anchor point for this loop is a H-bond between His26 and Pro44 which breaks often in the simulations (Figures 5 and S8). After this bond is broken, the 19-36 Ω loop can adopt many conformations. These conformations are mostly independent from the remaining structure as seen in Figure 5D where this loop is highly mobile during the run, while the 40-57 Ω loop remains nearly unchanged compared to its starting conformation, presumably because it is anchored to the heme by the hydrogen bond network discussed earlier.

**Figure 4.**
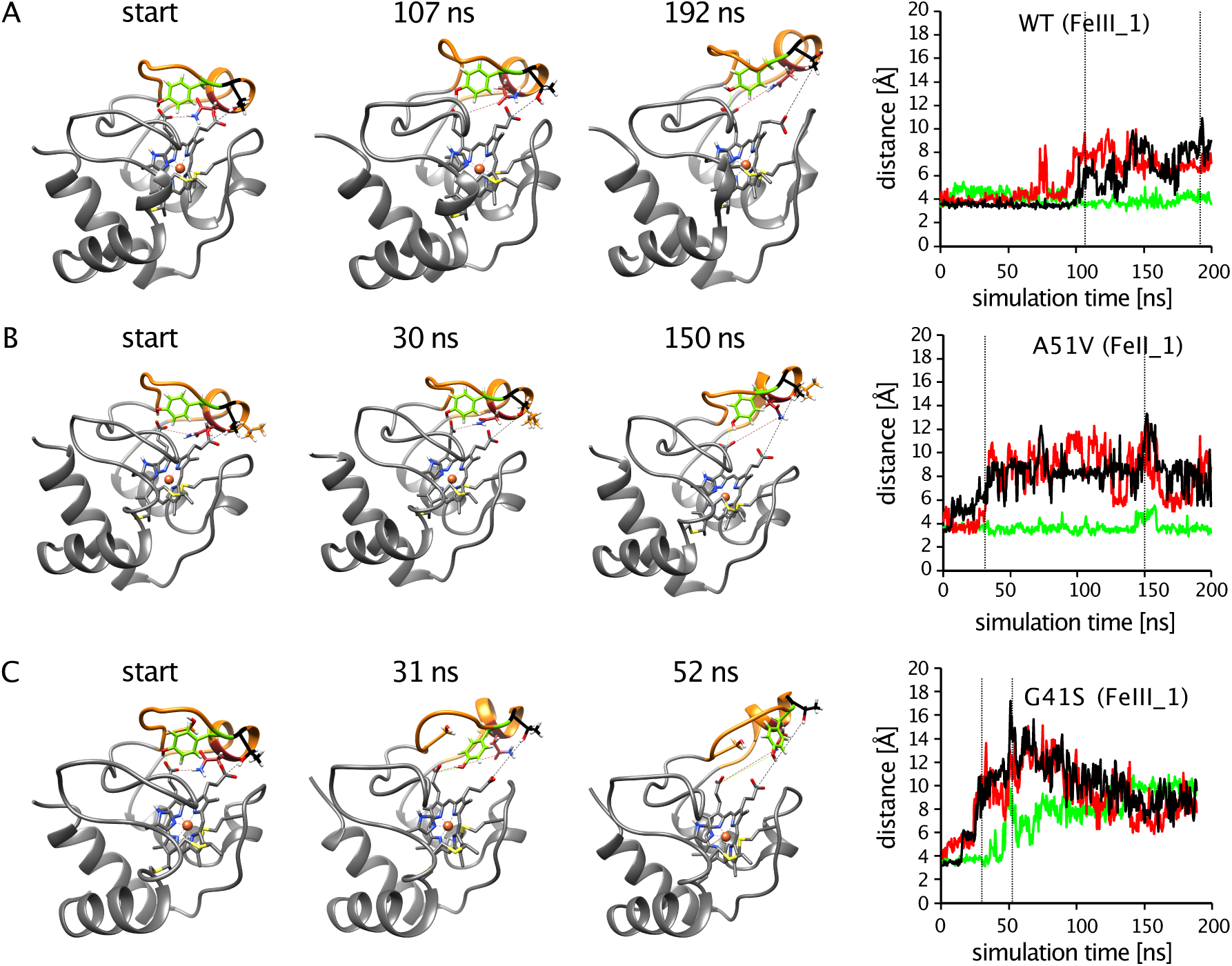
Molecular dynamics of the 40-57 Ω loop. (A-C) Selected snapshots and critical distances for WT (FeIII_1) (A), A51V (FeII_1) (B) and G41S (FeIII_1) (C). Residues shown are Tyr48 in light green, Thr49 in black and Asn52 in red (A-C); Val51 in orange (B), Ser41 in orange (C). The 40-57 Ω loop is in orange. Right hand panels show distances for Tyr48 hydroxyl to heme propionate (CGA) (light green), Thr49 hydroxyl to heme propionate (CGD) (black) and Asn52 amide (ND2) to heme propionate (CGA) (red) over the entire simulation. Dotted lines indicate times of structure snapshots. Distances for all simulations are in Figure S7.

**Figure 5.**
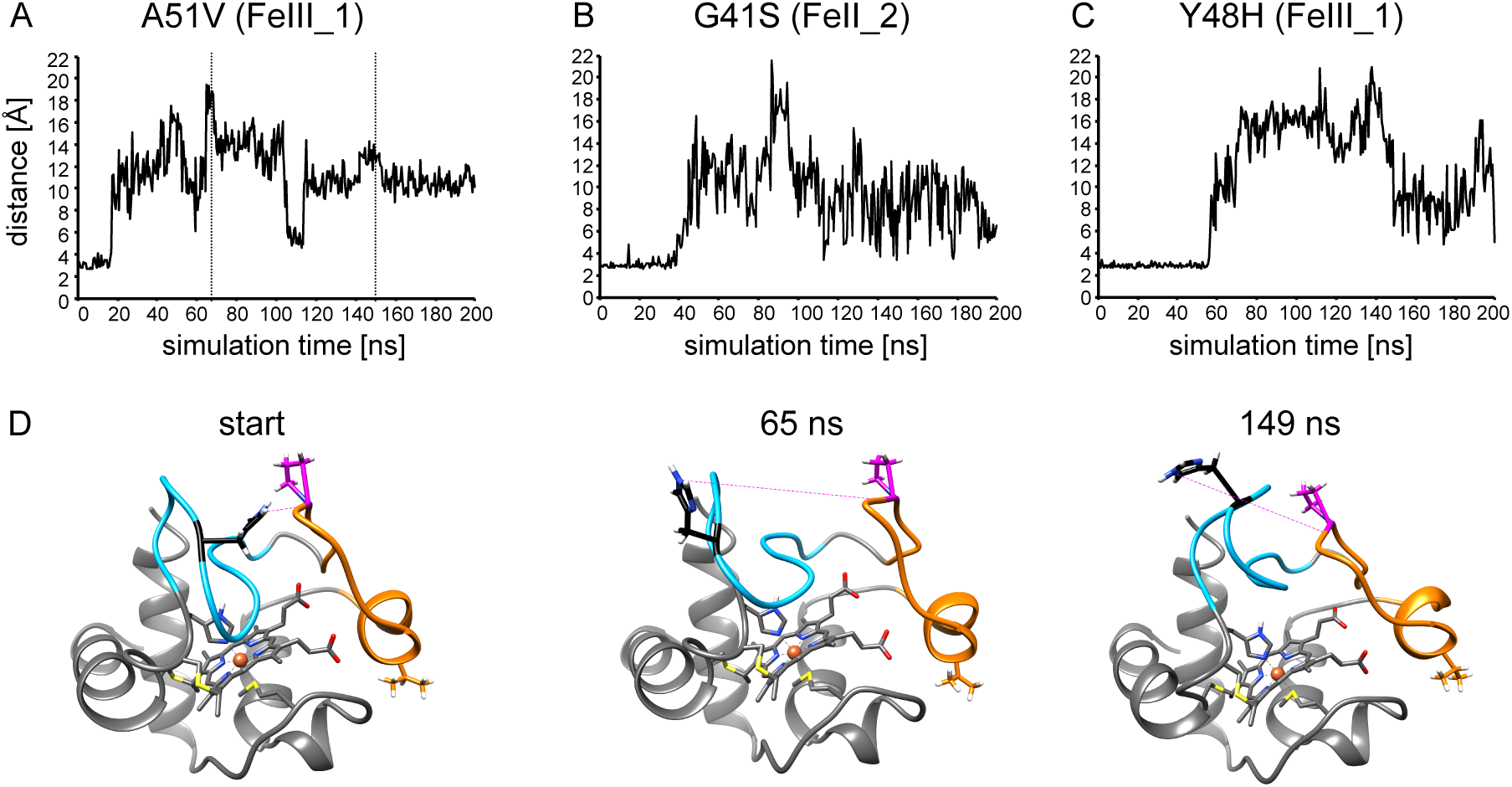
Molecular dynamics of the 19-36 Ω-loop. (A-C) His26 amine (NE2) to Pro44 carbonyl distance for A51V (FeIII_1) (A), G41S (FeII_2) (B) and Y48H (FeIII_1) (C). Dotted lines in (A) indicate times of structure snapshots. (D) Selected snapshots for A51V (FeIII_1) at the start, 65 ns and 149 ns. Resides shown are His26 in black, Pro44 in magenta, Val51 in orange. The 19-36 Ω loop is in cyan and the 40-57 Ω loop is in orange. Distances for all simulations are in Figure S8.

### Correlation of conformational mobility with cytochrome *c* activities

To determine whether altered main chain dynamics in the Ω loops may underly the observed changes in apoptosome activation and/or peroxidase activity we examined the correlation with average RMSF for each Ω loop (Figure 6). Increased apoptosome activation was associated with increased average RMSF of the loops, in particular the 40-57 and 19-36 Ω loops. In contrast the strongest association for peroxidase activity was a negative correlation with average RMSF of the 71-85 Ω loop. While the Pearson correlations were not significant (Table 2), the relationships may be non-linear, in which case the Pearson test is not informative.

**Table 2.** Analysis of relationship between Ω loop RMSF and cytochrome *c* activities.

|  | apoptosome activation (relative) |  |  | peroxidase activity (M<sup>-1</sup>s<sup>-1</sup>) |  |  |
| --- | --- | --- | --- | --- | --- | --- |
| | 19-36 $\Omega$ loop | 40-57 $\Omega$ loop | 71-85 $\Omega$ loop | 19-36 $\Omega$ loop | 40-57 $\Omega$ loop | 71-85 $\Omega$ loop |
| *r* | 0.73 | 0.78 | 0.61 | -0.21 | 0.03 | -0.80 |
| *P* | 0.10 | 0.07 | 0.20 | 0.73 | 0.97 | 0.11 |

Pearson correlation *r* and *P* value for average RMSF for Cα in each region from simulation data for either ferrous (apoptosome activation) or ferric (peroxidase activity) cytochromes *c*.

**Figure 6.**
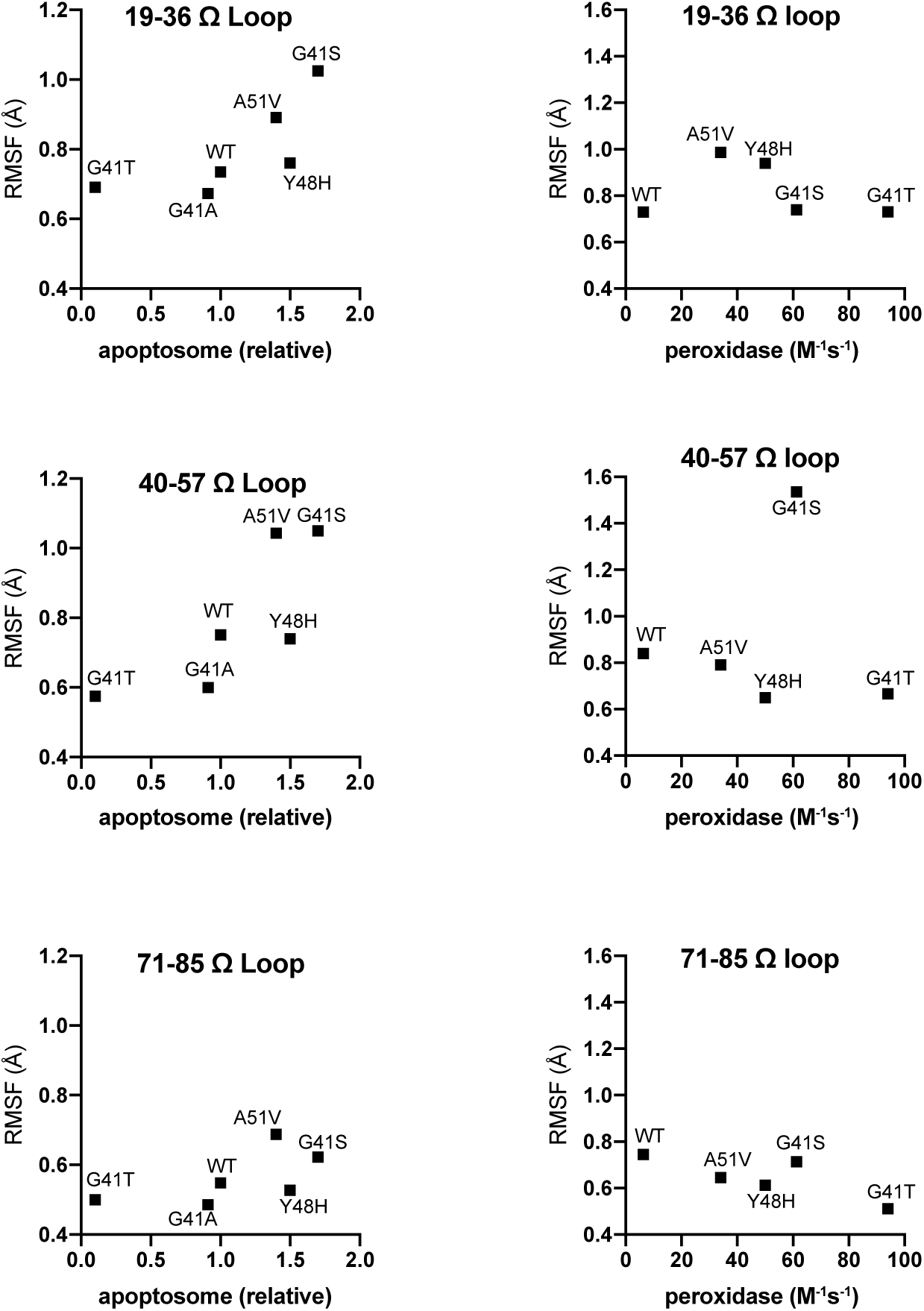
Correlation of apoptosome activation (left hand panels) or peroxidase activity (right hand panels) with RMSF for the 29-36 Ω loop, the 40-57 Ω loop, and the 71-85 Ω loop.

Peroxidase activity has previously been reported to be controlled by orientation of the Arg38 sidechain (27) and opening of specific cavities (37), via regulation of substrate access to the heme. From analysis of the crystal structures we hypothesized that the Arg38 sidechain may be mobile, and this is confirmed in the MD simulations. While the Arg38 mainchain shows very little movement, the side chain can adopt many conformations between three general states (Figures 7 and S9). The first orientation is modelled in the G41S crystal structure and serves as the starting point in all simulations; it is typified by a ∼15 Å distance from the guanidino carbon to the heme Fe, and can be relatively stable as seen in a G41A simulation (Figure 6A). The second orientation faces towards the heme Fe (∼11 Å distance), and is observed in the other crystal structures. Figure 6B shows a simulation with WT in which Arg38 adopts this heme facing conformation within 60 ns and remains in this conformation. The third conformation is not observed in crystal structures and faces towards the surface of the protein (∼17 Å distance to heme). This conformation can also be stable as seen in a G41S simulation (Figure 6C). In WT Arg38 mostly adopts the heme-facing conformation, consistent with the orientation in the WT structure, whereas for all variants Arg38 mainly populates a mix of the heme-and surface-facing conformations (Figure S9). Interestingly for G41S, Y48H, A51V and G41A cytochromes *c*, the surface-facing conformation is favored in the ferric state and the heme-facing conformation in the ferrous state.

**Figure 7.**
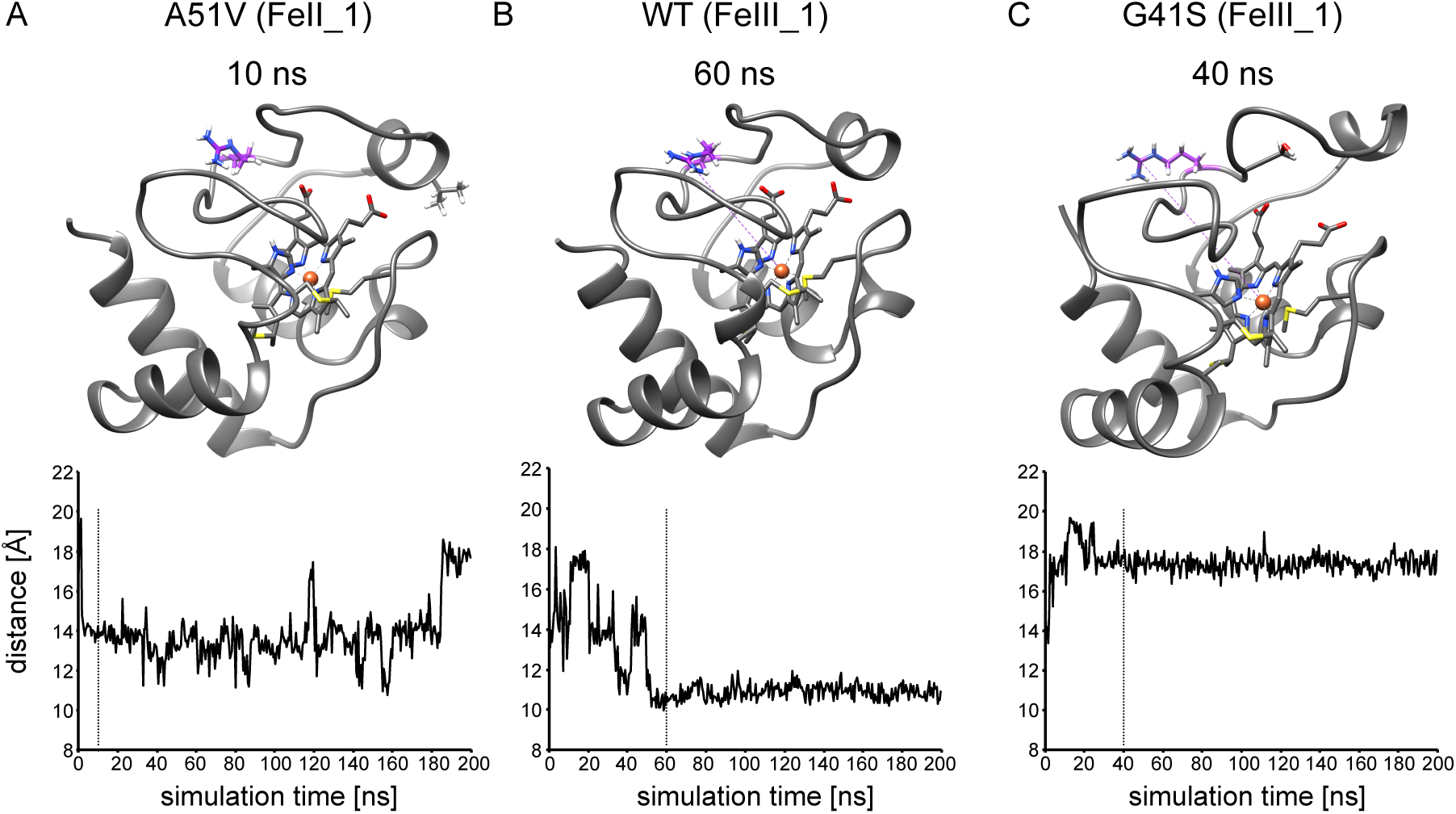
Molecular dynamics of Arg38 side chain. Selected snapshots and Arg38 guanidino carbon (CZ) to heme Fe distance for A51V (FeII_1) (A), WT (FeIII_1) (B) and G41S (FeIII_1) (C). Arg38 sidechain in purple. Dotted lines indicate times of structure snapshots. Distances for all simulations are in Figure S9.

To further investigate the role of the Arg38 sidechain in the peroxidase reaction, we expressed the R38W variant of cytochrome *c*. This variant was originally reported as a low frequency SNP (rs11548791, p.Arg39Trp, (38)). However it is now unclear if this is a real variant (39), and gnomAD v2.1.1 reports an alternative low frequency SNP (rs1047899818, p.Arg39Gln) (40). Nonetheless, as Trp has a larger van der Waals volume than Arg (0.79 nm^3^ compared to 0.59 nm^3^) and therefore might be expected to block access to substrate, we hypothesized that this variant would decrease the peroxidase activity of cytochrome *c*. The R38W cytochrome *c* variant was folded correctly with a hexacoordinate Fe, a slight blue shift of the Soret band, an unchanged *T*_*m*_, but a lowered redox potential when compared to WT (Table S1). Somewhat surprisingly peroxidase activity was increased to ∼60 M^-1^s^-1^ and showed a lag phase similar to other variants (Figure S10). This can be interpreted in two ways. The Arg38 sidechain may not control access of peroxidase substrates, and the R38W mutant alters peroxidase activity by another mechanism. Alternatively, rather than the indole sidechain assuming a WT-like position and blocking access to the peroxidase site, it could be rotated into solvent, exposing the water molecules and a substrate access channel otherwise sequestered behind the guanidino group of arginine.

The degree of opening of two cavities, A and B (defined by the Ala50 - Gly77 and Asn31-Ala43 Cα distances respectively), in yeast iso-1-cytochrome *c* has also been proposed to regulate substrate access to the heme (37). We therefore analyzed these distances to determine whether there was any difference in cavity opening between the variants (Figure S11). For cavity A (Figure S11A) there was very little variation in the Ala50-Gly77 distance apart from three simulations in which the distance increased coincident with unfolding of the 40-57 Ω loop described earlier. No simulations showed cavity A in the closed state as defined by Bortolotti *et al*. The distance of cavity B (Figure S11B) showed similar variation in all simulations, with no indication of more time spent in the open state in the variants with higher peroxidase activity.

Our simulations also provide an opportunity to test conclusions made from other MD data regarding the role of conformational dynamics in influencing the redox potential and the alkaline transition p*K*. Recently Oviedo-Rouco *et al* reported 50 ns simulations of equine ferric cytochrome *c* and selected variants (30). They concluded that a decrease in redox potential correlated with an increase in RMSF for the entire protein, the 70-85 Ω loop, and to a lesser extent the 40-57 Ω loop. The variants studied here have a similarly wide range of redox potential (189 - 244 mV). However, with our much more extensive MD data (400 ns for each ferric variant) there is no correlation between redox potential and RMSF (Figure S12, Table S3). In our variants with p*K* ranging from 6.7 - 9.3 there is an indication of the reported positive correlation between the alkaline transition p*K* and RMSF of the 70-85 Ω loop, however this is not significant (Table S4).

## Discussion

The ability of cytochrome *c* to contribute to multiple cellular processes is ascribed to its ability to switch between conformational states. By combining measurement of activities and properties of WT and five variant human cytochromes *c* with structural analyses and MD simulations we have been able to test existing theories of the role of conformational dynamics in controlling activities and properties of cytochrome *c*, and gain new insights into the potential importance of conformational dynamics in the pro-apoptotic activities of cytochrome *c*. The high structural similarity of cytochromes *c* with a wide range of redox potential, alkaline transition p*K*, peroxidase activity and apoptosome activation suggests that changes in these properties could be driven by variation in protein dynamics in local regions of cytochrome *c*.

The MD data for WT, G41S and Y41H cytochromes *c* are in general agreement with measurements of conformational dynamics by NMR-based hydrogen-deuterium exchange and ^15^N relaxation experiments (7, 28), and a previous report of 30 ns MD simulations (29), with increased movement for the pathogenic variants, particularly in the 19-36 and 40-57 Ω loops. This suggests that the MD data we have obtained provides a representative indication of the conformational dynamics of cytochromes *c*, and therefore is useful for assessing relationships between movement of various regions of cytochrome c and its properties and activities. Interestingly our data do not support the recent conclusion, from much shorter simulations of a range of equine cytochrome *c* variants, that redox potential is modulated by movement of the 40-57 and 71-85 Ω loops (30). Our data did provide partial support for the previous conclusion that flexibility of the 71-85 Ω may regulate the alkaline transition p*K* (30). The differences between our repeated, independent 200 ns simulations for each variant emphasizes that one should take care when making conclusions from short MD simulations of cytochrome *c*. Even our comparatively long simulations, although supporting conclusions from other methods, show significant stochastic variation.

Apoptosome formation requires the interaction of cytochrome *c* with Apaf-1. The primary driver of this interaction is formation of ionic bonds involving multiple lysine residues distributed across the surface of cytochrome *c* (41, 42). All the pathogenic variants showed a modest increase of in vitro apoptosome activation (maximum of ∼two-fold increase), whereas the G41A mutation has no impact and the G41T mutation causes an approximate ten-fold decrease. Increased RMSD correlates with increased apoptosome activation, however the mechanism underlying increased RMSD differs between the pathogenic variants. For G41S cytochrome *c* the main contributor is increased RMSF of residues in the 40-57 Ω loop, whereas for A51V and Y48H cytochromes *c* the major contributions are from the 19-36 Ω loop. The ground and active state cryo-EM apoptosome structures suggest that cytochrome *c* initially interacts with the WD2 domain of Apaf-1 (24, 25), with this interaction involving the 40-57 Ω loop. Subsequently there is a ∼60° rotation of the WD1 domain, locking cytochrome *c* in place. The increased flexibility caused by the pathogenic mutations may therefore facilitate binding by an induced fit mechanism, rather than by changing specific interactions once cytochrome *c* is bound.

To become an active peroxidase, cytochrome *c* must be converted from its native hexacoordinate state to a state in which the heme Fe is pentacoordinate. Our previous data shows that mutation of Gly41 to Ser or Thr increases peroxidase activity by facilitating the oxidation of the Met80 sulfur (18). This triggers conversion of the Fe to the active pentacoordinate state, and once in this state the cytochrome *c* variants have the same activity. A possible explanation for the increased activity of the mutations is that they enhance accessibility of the heme to peroxide, and previous studies have suggested that rigidity of the 71-85 Ω loop acts to inhibit peroxidase activity (43). Our extensive MD data with additional variants does not support a hypothesis that increased movement of the cytochrome *c* main chain, particularly in the 71-85 Ω loop, enhances conversion to the active state. In particular, G41T cytochrome *c*, the variant with highest peroxidase activity, has much lower overall flexibility than WT, and there was a negative correlation between peroxidase activity and RMSF of the 71-85 Ω loop. We also investigated alternative mechanisms for controlling peroxidase activity, orientation of the Arg38 sidechain (13,19,27), and the opening of channels providing access to the heme (37). However, neither of these mechanisms were supported by our data. An alternative impact of the mutations may be stabilization of a minor pentacoordinate Fe state (7), an event that will not be captured in our MD simulations where cytochrome *c* remains hexacoordinate.

Overall our data show that pathogenic variants in cytochrome *c* enhance the activities of cytochrome *c* that are associated with apoptosis. However, the current consensus is that apoptosis must be restrained to enable normal platelet production (44). Furthermore, peripheral blood mononuclear cells from subjects with G41S cytochrome *c* do not show altered susceptibility to apoptotic stimuli (12). Interestingly, in the context of mouse cytochrome *c*, the G41S mutation causes increased peroxidase activity, decreased apoptosome activation, and is not associated with thrombocytopenia (13, 17). This suggests that it may be the combination of impact on apoptosome activation and peroxidase activity that triggers abnormal platelet formation. The mechanism by which the pathogenic variants disrupt platelet formation therefore requires further investigation.

## Experimental Procedures

### Expression, purification and characterization of cytochrome c variants

The Y48H and A51V mutations were introduced into the human cytochrome *c* expression vector pBTR(HumanCc) (45) by site-directed mutagenesis, and the presence of the mutation was confirmed by DNA sequencing. WT, G41S, G41T, Y48H and A51V variants were expressed and purified as previously described (22), and the presence of each mutation was confirmed by MALDI analysis. The concentration of cytochrome *c* was measured using an extinction coefficient of 106.1 mM^-1^ cm^-1^ at 410 nm. Measurement of the redox potential and monitoring Met80 ligation by A_695_ was performed as described earlier (14). Global stability of ferric cytochrome *c* (40 μM in 50 mM citric acid buffer pH 3.8) was determined as previously described by differential scanning calorimetry (DSC) using a NanoDSC (TA instruments) (14) and was unchanged (Table S1).

### Apoptosome activation and peroxidase activity

Apoptosome activation was determined as previously described (14). Cytosolic extract was prepared from human monocytic lymphoma U937 cells and stored at -80 °C. Purified cytochrome *c* was added to cytosolic extract (100 µg protein) in the presence of 1 mM dATP in a final volume of 100 µL caspase assay buffer (100 mM HEPES– KOH (pH 7.25), 10% w/v sucrose, 0.1% w/v CHAPS, and 5 mM DTT) and incubated at 37 °C. Caspase 3-like activity was measured fluorometrically by the addition of 50 µM acetyl-Asp-Glu-Val-Asp-7-amino-4-methylcoumarin (Ac-DEVD-AMC) at 37 °C in 96-well plates using a ClarioSTAR microplate reader (BMG LABTECH). The production of AMC (λ_EM_ = 390 nm, λ_EX_= 460 nm) was quantified by reference to an AMC standard curve. Apoptosome activation was calculated from the maximum slope of a plot of the rate of AMC production versus time. The standard deviation error of the slope was calculated according to the method described elsewhere (46). All values were normalized to WT = 1.

The second order rate constants for the peroxidase activity of the cytochrome *c* variants were determined as previously described (18) using 3,5’,5,5’-tetramethylbenzidine (TMB). Cytochrome *c* (5 μM) was incubated in the presence of TMB (1.6 mM) and H_2_O_2_ (25-300 μM) in phosphate buffer (pH 5.4, 80 mM) at room temperature in a 96-well plate with a total volume of 100 μL. The absorption maximum at 652 nm and an extinction coefficient of 39,000 M^-1^cm^-1^ was used to follow time dependent TMB oxidation. The second order rate constants were determined from the plot of reaction velocity versus H_2_O_2_ concentration.

### Molecular Dynamics

All calculations were based on the G41S variant structure 3NWV chain A (22) which were modified in COOT (47) to obtain starting models for WT, G41A, G41S, G41T, Y48H and A51V. Every system was simulated in both oxidized heme (FeIII) form and the reduced heme (FeII) form. Structures were solvated with TIP3P water (48) in a box approximately 66 Å by 61 Å by 66 Å and the total charge neutralized with seven or eight chloride atoms. The protonation state for each model was set as appropriate at pH 7.0 (49) using the PDB2PQR webserver (50). VMD packages autoionize, solvate (51) and cytochrome *c* specific topology files (52) were used to prepare simulation systems. All B factors were set to 1. The simulations were performed at NeSI (NZ eScience Infrastructure). All simulations were done using NAMD version 2.11 (53), with the CHARMM36 force-field (48,54,55) and cytochrome c heme parameters taken from (52). All simulations were done with a timestep of 2 fs using the RATTLE (56) and SETTLE (57) algorithms. Simulated system data were saved every 10 ps. Simulation systems were equilibrated for 16 ns followed by productive MD run of 200 ns. Each system was run in duplicate. Simulations were performed using the NPT ensemble with the pressure set to 1 atmosphere and the temperature set to 298K. The pressure and temperature were maintained using the Langevin piston and the Langevin theromostat respectively. Electrostatics were calculated with the particle mesh Ewald method (58). The van der Waals interactions were truncated smoothly between 10 and 12 Å.

### Crystallization of Y48H and G41T cytochromes *c*

For the Y48H variant hanging drops of 1 μL of 29.5 mg/mL of reduced protein and 2 μL reservoir buffer were allowed to equilibrate above the reservoir buffer (32% (w/v) polyethylene glycol 5000, 50 mM lithium sulfate, 50 mM Tris-HCl pH 8.5). The protein solution was 17.5 mM sodium phosphate, 25 mM sodium chloride, 15 mM sodium dithionite (pH 7.6). For the G41T variant hanging drops of 1 μL of 30.1 mg/mL of reduced protein and 1 μL reservoir buffer were allowed to equilibrate above the reservoir buffer (30% (w/v) polyethylene glycol 5000, 60 mM lithium sulfate, 50 mM Tris-HCl pH 8.5). The protein solution was 26 mM sodium phosphate, 37 mM sodium chloride, 14 mM sodium dithionite (pH 7.6). In both cases the sodium dithionite was freshly prepared to ensure complete reduction of the protein. For both variants, crystals appeared at 18 °C after a day as long thin red needles up to 100 µm in length. Both crystals were harvested after 5 days. In general, all needles appeared to have a very short window of high-quality diffraction, with needles harvested too early or too late often showing no diffraction. Further details including data collection, reduction and refinements statistics are in supplementary information.

### Statistical analyses

All statistical analyses were done in GraphPad Prism v8.

## Data availability

The datasets generated and analyzed during the current study are available in the Protein Data Bank (PDB) with the following codes: 6ECJ (G41T human cytochrome *c* variant) and 5EXQ (Y48H human cytochrome *c* variant).

## Acknowledgements

The authors wish to acknowledge the contribution of NeSI high-performance computing facilities to the results of this research. NZ’s national facilities are provided by the NZ eScience Infrastructure and funded jointly by NeSI’s collaborator institutions and through the Ministry of Business, Innovation & Employment’s Research Infrastructure programme (https://www.nesi.org.nz). We would like to thank Dr Matteo Ferla for assistance with molecular simulations.

## Abbreviations

Apaf-1: apoptotic protease activating factor-1
H_2_O_2_: hydrogen peroxide
H-bond: hydrogen bond
hp: heme propionate
MD: molecular dynamics
RMSD: root mean square deviation
RMSF: root mean square fluctuation
*T*_*m*_: melting temperature
TMB: 3,5’,5,5’-tetramethylbenzidine

## References

1. Alvarez-Paggi, D., Hannibal, L., Castro, M. A., Oviedo-Rouco, S., Demicheli, V., Tortora, V., Tomasina, F., Radi, R., and Murgida, D. H. (2017) Multifunctional Cytochrome c: Learning New Tricks from an Old Dog. Chem. Rev. 117, 13382–13460

2. Hüttemann, M., Pecina, P., Rainbolt, M., Sanderson, T. H., Kagan, V. E., Samavati, L., Doan, J. W., and Lee, I. (2011) The multiple functions of cytochrome c and their regulation in life and death decisions of the mammalian cell: From respiration to apoptosis. Mitochondrion 11, 369–381

3. Li, K., Li, Y., Shelton, J. M., Richardson, J. A., Spencer, E., Chen, Z. J., Wang, X., and Williams, R. S. (2000) Cytochrome c deficiency causes embryonic lethality and attenuates stress-induced apoptosis. Cell 101, 389–399

4. Hao, Z., Duncan, G. S., Chang, C. C., Elia, A., Fang, M., Wakeham, A., Okada, H., Calzascia, T., Jang, Y., You-Ten, A., Yeh, W.-C., Ohashi, P., Wang, X., and Mak, T. W. (2005) Specific ablation of the apoptotic functions of cytochrome c reveals a differential requirement for cytochrome c and Apaf-1 in apoptosis. Cell 121, 579–591

5. Kagan, V. E., Bayir, H. A., Belikova, N. A., Kapralov, O., Tyurina, Y. Y., Tyurin, V. A., Jiang, J., Stoyanovsky, D. A., Wipf, P., Kochanek, P. M., Greenberger, J. S., Pitt, B., Shvedova, A. A., and Borisenko, G. (2009) Cytochrome c/cardiolipin relations in mitochondria: a kiss of death. Free Radic. Biol. Med. 46, 1439–1453

6. Hannibal, L., Tomasina, F., Capdevila, D. A., Demicheli, V., Tórtora, V., Alvarez-Paggi, D., Jemmerson, R., Murgida, D. H., and Radi, R. (2016) Alternative Conformations of Cytochrome c: Structure, Function, and Detection. Biochemistry 55, 407–428

7. Deacon, O. M., Karsisiotis, A. I., Moreno-Chicano, T., Hough, M. A., Macdonald, C., Blumenschein, T. M. A., Wilson, M. T., Moore, G. R., and Worrall, J. A. R. (2017) Heightened Dynamics of the Oxidized Y48H Variant of Human Cytochrome c Increases Its Peroxidatic Activity. Biochemistry 56, 6111–6124

8. Morison, I. M., Cramer Borde, E. M., Cheesman, E. J., Cheong, P. L., Holyoake, A. J., Fichelson, S., Weeks, R. J., Lo, A., Davies, S. M., Wilbanks, S. M., Fagerlund, R. D., Ludgate, M. W., da Silva Tatley, F. M., Coker, M. S., Bockett, N. A., Hughes, G., Pippig, D. A., Smith, M. P., Capron, C., and Ledgerwood, E. C. (2008) A mutation of human cytochrome c enhances the intrinsic apoptotic pathway but causes only thrombocytopenia. Nat. Genet. 40, 387–389

9. De Rocco, D., Cerqua, C., Goffrini, P., Russo, G., Pastore, A., Meloni, F., Nicchia, E., Moraes, C. T., Pecci, A., Salviati, L., and Savoia, A. (2014) Mutations of cytochrome c identified in patients with thrombocytopenia THC4 affect both apoptosis and cellular bioenergetics. Biochim. Biophys. Acta 1842, 269–274

10. Johnson, B., Lowe, G. C., Futterer, J., Lordkipanidzé, M., MacDonald, D., Simpson, M. A., Sanchez Guiu, I., Drake, S., Bem, D., Leo, V., Fletcher, S. J., Dawood, B., Rivera, J., Allsup, D., Biss, T., Bolton-Maggs, P. H., Collins, P., Curry, N., Grimley, C., James, B., Makris, M., Motwani, J., Pavord, S., Talks, K., Thachil, J., Wilde, J., Williams, M., Harrison, P., Gissen, P., Mundell, S., Mumford, A., Daly, M. E., Watson, S. P., Morgan, N. V., and Group, U. G. S. (2016) Whole exome sequencing identifies genetic variants in inherited thrombocytopenia with secondary qualitative function defects. Haematologica 101, 1170–1179

11. Uchiyama, Y., Yanagisawa, K., Kunishima, S., Shiina, M., Ogawa, Y., Nakashima, M., Hirato, J., Imagawa, E., Fujita, A., Hamanaka, K., Miyatake, S., Mitsuhashi, S., Takata, A., Miyake, N., Ogata, K., Handa, H., Matsumoto, N., and Mizuguchi, T. (2018) A novel CYCS mutation in the α-helix of the CYCS C-terminal domain causes non-syndromic thrombocytopenia. Clin. Genet. 94, 548–553

12. Ong, L., Morison, I. M., and Ledgerwood, E. C. (2017) Megakaryocytes from CYCS mutation-associated thrombocytopenia release platelets by both proplatelet-dependent and -independent processes. Br. J. Haematol. 176, 268–279

13. Josephs, T. M., Morison, I. M., Day, C. L., Wilbanks, S. M., and Ledgerwood, E. C. (2014) Enhancing the peroxidase activity of cytochrome c by mutation of residue 41: implications for the peroxidase mechanism and cytochrome c release. Biochem. J. 458, 259–265

14. Josephs, T. M., Liptak, M. D., Hughes, G., Lo, A., Smith, R. M., Wilbanks, S. M., Bren, K. L., and Ledgerwood, E. C. (2013) Conformational change and human cytochrome c function: mutation of residue 41 modulates caspase activation and destabilizes Met-80 coordination. J. Biol. Inorg. Chem. 18, 289–297

15. Deacon, O. M., White, R. W., Moore, G. R., Wilson, M. T., and Worrall, J. A. R. (2019) Comparison of the structural dynamic and mitochondrial electron-transfer properties of the proapoptotic human cytochrome c variants, G41S, Y48H and A51V. J. Inorg. Biochem. 203, 110924

16. Lei, H., and Bowler, B. E. (2019) Naturally Occurring A51V Variant of Human Cytochrome c Destabilizes the Native State and Enhances Peroxidase Activity. J. Phys. Chem. B 123, 8939–8953

17. Josephs, T. M., Hibbs, M. E., Ong, L., Morison, I. M., and Ledgerwood, E. C. (2015) Interspecies Variation in the Functional Consequences of Mutation of Cytochrome c. PLoS One 10, e0130292

18. Parakra, R. D., Kleffmann, T., Jameson, G. N. L., and Ledgerwood, E. C. (2018) The proportion of Met80-sulfoxide dictates peroxidase activity of human cytochrome c. Dalton Trans 47, 9128–9135

19. Rajagopal, B. S., Edzuma, A. N., Hough, M. A., Blundell, K. L., Kagan, V. E., Kapralov, A. A., Fraser, L. A., Butt, J. N., Silkstone, G. G., Wilson, M. T., Svistunenko, D. A., and Worrall, J. A. (2013) The hydrogen-peroxide-induced radical behaviour in human cytochrome c-phospholipid complexes: implications for the enhanced pro-apoptotic activity of the G41S mutant. Biochem. J. 456, 441–452

20. Ledgerwood, E. C., Dunstan-Harrison, C., Ong, L., and Morison, I. M. (2019) CYCS gene variants associated with thrombocytopenia. Platelets 30, 672–674

21. Mugnol, K. C., Ando, R. A., Nagayasu, R. Y., Faljoni-Alario, A., Brochsztain, S., Santos, P. S., Nascimento, O. R., and Nantes, I. L. (2008) Spectroscopic, structural, and functional characterization of the alternative low-spin state of horse heart cytochrome C. Biophys. J. 94, 4066–4077

22. Liptak, M. D., Fagerlund, R. D., Ledgerwood, E. C., Wilbanks, S. M., and Bren, K. L. (2011) The proapoptotic G41S mutation to human cytochrome c alters the heme electronic structure and increases the electron self-exchange rate. J. Am. Chem. Soc. 133, 1153–1155

23. Krishna, M. M., Lin, Y., Rumbley, J. N., and Englander, S. W. (2003) Cooperative omega loops in cytochrome c: role in folding and function. J. Mol. Biol. 331, 29–36

24. Zhou, M., Li, Y., Hu, Q., Bai, X. C., Huang, W., Yan, C., Scheres, S. H., and Shi, Y. (2015) Atomic structure of the apoptosome: mechanism of cytochrome c- and dATP-mediated activation of Apaf-1. Genes Dev. 29, 2349–2361

25. Cheng, T. C., Hong, C., Akey, I. V., Yuan, S., and Akey, C. W. (2016) A near atomic structure of the active human apoptosome. Elife 5, e17755

26. Li, Y., Zhou, M., Hu, Q., Bai, X.-c., Huang, W., Scheres, S. H. W., and Shi, Y. (2017) Mechanistic insights into caspase-9 activation by the structure of the apoptosome holoenzyme. Proc. Natl. Acad. Sci. U. S. A. 114, 1542–1547

27. McClelland, L. J., Mou, T.-C., Jeakins-Cooley, M. E., Sprang, S. R., and Bowler, B. E. (2014) Structure of a mitochondrial cytochrome c conformer competent for peroxidase activity. Proc. Natl. Acad. Sci. U. S. A. 111, 6648–6653

28. Karsisiotis, A. I., Deacon, O. M., Wilson, M. T., Macdonald, C., Blumenschein, T. M., Moore, G. R., and Worrall, J. A. (2016) Increased dynamics in the 40-57 Omega-loop of the G41S variant of human cytochrome c promote its pro-apoptotic conformation. Sci. Rep. 6, 30447

29. Muneeswaran, G., Kartheeswaran, S., Pandiaraj, M., Muthukumar, K., Sankaralingam, M., and Arunachalam, S. (2017) Investigation of structural dynamics of Thrombocytopenia Cargeeg mutants of human apoptotic cytochrome c: A molecular dynamics simulation approach. Biophys. Chem. 230, 117–126

30. Oviedo-Rouco, S., Perez-Bertoldi, J. M., Spedalieri, C., Castro, M. A., Tomasina, F., Tortora, V., Radi, R., and Murgida, D. H. (2020) Electron transfer and conformational transitions of cytochrome c are modulated by the same dynamical features. Arch. Biochem. Biophys. 680, 108243

31. Oviedo-Rouco, S., Castro, M. A., Alvarez-Paggi, D., Spedalieri, C., Tortora, V., Tomasina, F., Radi, R., and Murgida, D. H. (2019) The alkaline transition of cytochrome c revisited: Effects of electrostatic interactions and tyrosine nitration on the reaction dynamics. Arch. Biochem. Biophys. 665, 96–106

32. Sargsyan, K., Grauffel, C., and Lim, C. (2017) How Molecular Size Impacts RMSD Applications in Molecular Dynamics Simulations. J. Chem. Theory Comput. 13, 1518–1524

33. Sawle, L., and Ghosh, K. (2016) Convergence of Molecular Dynamics Simulation of Protein Native States: Feasibility vs Self-Consistency Dilemma. J. Chem. Theory Comput. 12, 861–869

34. Garcia-Heredia, J. M., Diaz-Moreno, I., Nieto, P. M., Orzaez, M., Kocanis, S., Teixeira, M., Perez-Paya, E., Diaz-Quintana, A., and De la Rosa, M. A. (2010) Nitration of tyrosine 74 prevents human cytochrome c to play a key role in apoptosis signaling by blocking caspase-9 activation. Biochim. Biophys. Acta 1797, 981–993

35. Garcia, A. E., and Hummer, G. (1999) Conformational dynamics of cytochrome c: correlation to hydrogen exchange. Proteins 36, 175–191

36. Muneeswaran, G., Kartheeswaran, S., Muthukumar, K., and Karunakaran, C. (2018) Temperature-dependent conformational dynamics of cytochrome c: Implications in apoptosis. J. Mol. Graph. Model. 79, 140–148

37. Bortolotti, C. A., Amadei, A., Aschi, M., Borsari, M., Corni, S., Sola, M., and Daidone, I. (2012) The reversible opening of water channels in cytochrome c modulates the heme iron reduction potential. J. Am. Chem. Soc. 134, 13670–13678

38. Lek, M., Karczewski, K. J., Minikel, E. V., Samocha, K. E., Banks, E., Fennell, T., O’Donnell-Luria, A. H., Ware, J. S., Hill, A. J., Cummings, B. B., Tukiainen, T., Birnbaum, D. P., Kosmicki, J. A., Duncan, L. E., Estrada, K., Zhao, F., Zou, J., Pierce-Hoffman, E., Berghout, J., Cooper, D. N., Deflaux, N., DePristo, M., Do, R., Flannick, J., Fromer, M., Gauthier, L., Goldstein, J., Gupta, N., Howrigan, D., Kiezun, A., Kurki, M. I., Moonshine, A. L., Natarajan, P., Orozco, L., Peloso, G. M., Poplin, R., Rivas, M. A., Ruano-Rubio, V., Rose, S. A., Ruderfer, D. M., Shakir, K., Stenson, P. D., Stevens, C., Thomas, B. P., Tiao, G., Tusie-Luna, M. T., Weisburd, B., Won, H. H., Yu, D., Altshuler, D. M., Ardissino, D., Boehnke, M., Danesh, J., Donnelly, S., Elosua, R., Florez, J. C., Gabriel, S. B., Getz, G., Glatt, S. J., Hultman, C. M., Kathiresan, S., Laakso, M., McCarroll, S., McCarthy, M. I., McGovern, D., McPherson, R., Neale, B. M., Palotie, A., Purcell, S. M., Saleheen, D., Scharf, J. M., Sklar, P., Sullivan, P. F., Tuomilehto, J., Tsuang, M. T., Watkins, H. C., Wilson, J. G., Daly, M. J., MacArthur, D. G., and Exome Aggregation, C. (2016) Analysis of protein-coding genetic variation in 60,706 humans. Nature 536, 285–291

39. Bertini, I., Grassi, E., Luchinat, C., Quattrone, A., and Saccenti, E. (2006) Monomorphism of human cytochrome c. Genomics 88, 669–672

40. Karczewski, K. J., Francioli, L. C., Tiao, G., Cummings, B. B., Alföldi, J., Wang, Q., Collins, R. L., Laricchia, K. M., Ganna, A., Birnbaum, D. P., Gauthier, L. D., Brand, H., Solomonson, M., Watts, N. A., Rhodes, D., Singer-Berk, M., Seaby, E. G., Kosmicki, J. A., Walters, R. K., Tashman, K., Farjoun, Y., Banks, E., Poterba, T., Wang, A., Seed, C., Whiffin, N., Chong, J. X., Samocha, K. E., Pierce-Hoffman, E., Zappala, Z., O’Donnell-Luria, A. H., Minikel, E. V., Weisburd, B., Lek, M., Ware, J. S., Vittal, C., Armean, I. M., Bergelson, L., Cibulskis, K., Connolly, K. M., Covarrubias, M., Donnelly, S., Ferriera, S., Gabriel, S., Gentry, J., Gupta, N., Jeandet, T., Kaplan, D., Llanwarne, C., Munshi, R., Novod, S., Petrillo, N., Roazen, D., Ruano-Rubio, V., Saltzman, A., Schleicher, M., Soto, J., Tibbetts, K., Tolonen, C., Wade, G., Talkowski, M. E., Neale, B. M., Daly, M. J., and MacArthur, D. G. (2019) The mutational constraint spectrum quantified from variation in 141,456 humans. BioRxiv

41. Yu, T., Wang, X., Purring-Koch, C., Wei, Y., and McLendon, G. L. (2001) A mutational epitope for cytochrome C binding to the apoptosis protease activation factor-1. J. Biol. Chem. 276, 13034–13038

42. Purring-Koch, C., and McLendon, G. (2000) Cytochrome c binding to Apaf-1: the effects of dATP and ionic strength. Proc. Natl. Acad. Sci. U. S. A. 97, 11928–11931

43. Nold, S. M., Lei, H., Mou, T.-C., and Bowler, B. E. (2017) Effect of a K72A Mutation on the Structure, Stability, Dynamics, and Peroxidase Activity of Human Cytochrome c. Biochemistry 56, 3358–3368

44. McArthur, K., Chappaz, S., and Kile, B. T. (2018) Apoptosis in megakaryocytes and platelets: the life and death of a lineage. Blood 131, 605–610

45. Olteanu, A., Patel, C. N., Dedmon, M. M., Kennedy, S., Linhoff, M. W., Minder, C. M., Potts, P. R., Deshmukh, M., and Pielak, G. J. (2003) Stability and apoptotic activity of recombinant human cytochrome c. Biochem. Biophys. Res. Commun. 312, 733–740

46. Taylor, J. (1982) An Introduction to Error Analysis, University Science Books. Mill Valley, California

47. Emsley, P., Lohkamp, B., Scott, W. G., and Cowtan, K. (2010) Features and development of Coot. Acta Crystallogr. D Biol. Crystallogr. 66, 486–501

48. MacKerell, A. D., Bashford, D., Bellott, M., Dunbrack, R. L., Evanseck, J. D., Field, M. J., Fischer, S., Gao, J., Guo, H., Ha, S., Joseph-McCarthy, D., Kuchnir, L., Kuczera, K., Lau, F. T., Mattos, C., Michnick, S., Ngo, T., Nguyen, D. T., Prodhom, B., Reiher, W. E., Roux, B., Schlenkrich, M., Smith, J. C., Stote, R., Straub, J., Watanabe, M., Wiorkiewicz-Kuczera, J., Yin, D., and Karplus, M. (1998) All-atom empirical potential for molecular modeling and dynamics studies of proteins. J. Phys. Chem. B 102, 3586–3616

49. Sondergaard, C. R., Olsson, M. H., Rostkowski, M., and Jensen, J. H. (2011) Improved Treatment of Ligands and Coupling Effects in Empirical Calculation and Rationalization of pKa Values. J. Chem. Theory Comput. 7, 2284–2295

50. Dolinsky, T. J., Nielsen, J. E., McCammon, J. A., and Baker, N. A. (2004) PDB2PQR: an automated pipeline for the setup of Poisson-Boltzmann electrostatics calculations. Nucleic Acids Res. 32, W665–667

51. Humphrey, W., Dalke, A., and Schulten, K. (1996) VMD: visual molecular dynamics. J. Mol. Graph. 14, 33-38, 27-38

52. Autenrieth, F., Tajkhorshid, E., Baudry, J., and Luthey-Schulten, Z. (2004) Classical force field parameters for the heme prosthetic group of cytochrome c. J. Comput. Chem. 25, 1613–1622

53. Phillips, J. C., Braun, R., Wang, W., Gumbart, J., Tajkhorshid, E., Villa, E., Chipot, C., Skeel, R. D., Kale, L., and Schulten, K. (2005) Scalable molecular dynamics with NAMD. J. Comput. Chem. 26, 1781–1802

54. Best, R. B., Zhu, X., Shim, J., Lopes, P. E., Mittal, J., Feig, M., and Mackerell, A. D., Jr. (2012) Optimization of the additive CHARMM all-atom protein force field targeting improved sampling of the backbone phi, psi and side-chain chi(1) and chi(2) dihedral angles. J. Chem. Theory Comput. 8, 3257–3273

55. MacKerell, A. D., Jr., Feig, M., and Brooks, C. L., 3rd. (2004) Improved treatment of the protein backbone in empirical force fields. J. Am. Chem. Soc. 126, 698–699

56. Andersen, H. C. (1983) Rattle - a Velocity Version of the Shake Algorithm for Molecular-Dynamics Calculations. J Comput Phys 52, 24–34

57. Miyamoto, S., and Kollman, P. A. (1992) Settle - an Analytical Version of the Shake and Rattle Algorithm for Rigid Water Models. J. Comput. Chem. 13, 952–962

58. Darden, T., York, D., and Pedersen, L. (1993) Particle Mesh Ewald - an N.Log(N) Method for Ewald Sums in Large Systems. J. Chem. Phys. 98, 10089–10092

