## Supplementary Information for "Altered structure and dynamics of pathogenic cytochrome c variants correlate with increased apoptotic activity"

**Contents:**

S-2 Crystallization of Y48H and G41T cytochromes *c*
S-3 Support information references
S-4 Table S1. Biophysical properties of cytochrome *c* variants
S-5 Table S2. X-ray data collection, reduction and refinement statistics
S-6 Table S3. RMSD for all MD simulations
S-7 Table S4. Correlation of RMSF with redox potential and alkaline transition p*K*
S-8 Figure S1. Structure and sequence of human ferric cytochrome *c*
S-9 Figure S2. Comparison of variant human cytochrome *c* structures
S-10 Figure S3. Comparison of the H-bond network of the human cytochrome *c* heme and 40-57 Ω-loop
S-11 Figure S4. Location of cytochrome *c* between the WD1 and WD2 domains of Apaf-1
S-12 Figure S5. RMSD histograms for combined MD simulations for WT cytochrome *c* and each variant
S-13 Figure S6. RMSF of Cα atoms as a function of amino acid in all simulations
S-14 Figure S7. Distances from Thr49, Asn52 and Tyr48/His48 to heme in all simulations
S-16 Figure S8. Distance from His26 to Pro44 in all simulations
S-17 Figure S9. Distance from Arg38 to the heme Fe in all simulations
S-18 Figure S10. Distances and RMSF for residues defining cavities A and B
S-19 Figure S11. Peroxidase activity of R38W cytochrome *c*
S-20 Figure S12. Correlation of RMSF with redox potential and alkaline transition p*K*

**Crystallization of Y48H and G41T cytochromes *c***

Our group was the first to report crystallization conditions for human cytochrome *c* (G41S) by a combination of dialysis and hanging drop format (1,2) utilizing a low ionic strength crystallization buffer with PEG at pH 7.5. Subsequently, these conditions have also been later utilized by Rajagopal *et al* for two additional human cytochrome *c* structures (3). While a large variety of crystallization conditions for eukaryotic cytochrome *c* can be found in the protein database, our continuing efforts in crystallizing human cytochrome *c* and its variants showed that nucleation is slow and very selective. Crystallization via dialysis appeared to be suboptimal and we investigated reported conditions as well as sparse matrix screening for new conditions, leading to a new ionic strength setup with PEG at pH 8.5.

Interestingly we found that Y48H crystallized in two crystal forms, as indistinguishable long red needles, both diffracting up to ~1.3 Å in either space group P1 or $P12_11$. Crystals in P1 appeared to be the same form as reported by Rajagopal *et al* (3) while crystals in $P12_11$ with two monomers in the asymmetric unit resembled the Y48H structure by Deacon *et al* (4) deposited after us. As the $P12_11$ dataset refined better, it was selected as our final model presented here.

G41T also crystallized as the same red needles, however in space group $P22_12_1$ with eight monomers in the asymmetric unit and never diffracted better than ~2.7 Å. We have described the G41T crystal form previously for our first G41S structure (2). The eight monomers are very similar with Cα RMSD of ~0.15-0.35 Å with chain C showing the overall lowest Bfactor of 17.5 $Å^2$. Chain H showed a poor density fit compared to the other chains, especially for surface loops, also resulting in the highest RMSD compared to the other chains. It was refined to 69 % chain occupancy with an overall B factor of 72.1 $Å^2$.

Data were collected at the MX2 beamline of the Australian Synchrotron at 93 K at 13.000 eV. For the Y48H crystal, two datasets were collected at different ends of the crystal needle and later merged. For the first dataset 266 frames at 0.5 degree at 180 mm detector distance were collected with 75% attenuation at 1 second exposure. For the second dataset 312 frames at 0.5 degree were collected with the detector distance being 350 mm and a 20 µm microcollimeter was used with 0% attenuation at 1 second exposure. For the G41T crystal, 180 frames at 1 degree at 400 mm detector distance were collected and a 20 µm microcollimeter was used with 0% attenuation at 0.7 second exposure. The Y48H data were indexed in space group $P12_11$ with unit cell dimensions of 57, 36 and 60 Å while the G41T data were indexed in space group $P2_12_11$ with unit cell dimensions of 65, 76 and 233 Å. All datasets were integrated in iMosflm (5). The two Y48H datasets were merged using the ccp4 program blend (6). Scaling was performed using aimless (7). Five % of reflections were reserved for the calculation of $R_{free}$ (Table S1). The data was cut to suggested resolution limits (Table S1) based on $CC_{1/2}$ and I/σ reported by aimless (7), extending the resolution further indicated problems with completeness for Y48H and with $R_{merge}$ for G41T. For the Y48H crystal chain A of human WT cytochrome *c* (PDB ID 3ZCF) (3) was used for molecular replacement in Phenix-Phaser (8) with two monomers in the asymmetric unit while for the G41T all eight chains of our first G41S crystal structure (2) with eight monomers in the asymmetric unit was used. Phenix-Refine (9) and COOT (10) were used for refinement. Coordinates and structure factors were deposited in the Protein Data Base as entry 5EXQ for Y48H and 6ECJ for G41T. Visualization, structure analysis and comparisons were performed using UCSF Chimera (11).

**Supporting Information References**


1. Liptak, M. D., Fagerlund, R. D., Ledgerwood, E. C., Wilbanks, S. M., and Bren, K. L. (2011) The proapoptotic G41S mutation to human cytochrome c alters the heme electronic structure and increases the electron self-exchange rate. *J. Am. Chem. Soc.* **133**, 1153-1155
2. Morison, I. M., Cramer Borde, E. M., Cheesman, E. J., Cheong, P. L., Holyoake, A. J., Fichelson, S., Weeks, R. J., Lo, A., Davies, S. M., Wilbanks, S. M., Fagerlund, R. D., Ludgate, M. W., da Silva Tatley, F. M., Coker, M. S., Bockett, N. A., Hughes, G., Pippig, D. A., Smith, M. P., Capron, C., and Ledgerwood, E. C. (2008) A mutation of human cytochrome c enhances the intrinsic apoptotic pathway but causes only thrombocytopenia. *Nat. Genet.* **40**, 387-389
3. Rajagopal, B. S., Edzuma, A. N., Hough, M. A., Blundell, K. L. I. M., Kagan, V. E., Kapralov, A. A., Fraser, L. A., Butt, J. N., Silkstone, G. G., Wilson, M. T., Svistunenko, D. A., and Worrall, J. A. R. (2013) The hydrogen-peroxide-induced radical behaviour in human cytochrome c-phospholipid complexes: implications for the enhanced pro-apoptotic activity of the G41S mutant. *Biochem. J.* **456**, 441-452
4. Deacon, O. M., Karsisiotis, A. I., Moreno-Chicano, T., Hough, M. A., Macdonald, C., Blumenschein, T. M. A., Wilson, M. T., Moore, G. R., and Worrall, J. A. R. (2017) Heightened Dynamics of the Oxidized Y48H Variant of Human Cytochrome c Increases Its Peroxidatic Activity. *Biochemistry* **56**, 6111-6124
5. Battye, T. G., Kontogiannis, L., Johnson, O., Powell, H. R., and Leslie, A. G. (2011) iMOSFLM: a new graphical interface for diffraction-image processing with MOSFLM. *Acta Crystallogr. D Biol. Crystallogr.* **67**, 271-281
6. Foadi, J., Aller, P., Alguel, Y., Cameron, A., Axford, D., Owen, R. L., Armour, W., Waterman, D. G., Iwata, S., and Evans, G. (2013) Clustering procedures for the optimal selection of data sets from multiple crystals in macromolecular crystallography. *Acta Crystallogr. D Biol. Crystallogr.* **69**, 1617-1632
7. Evans, P. R., and Murshudov, G. N. (2013) How good are my data and what is the resolution? *Acta Crystallogr. D Biol. Crystallogr.* **69**, 1204-1214
8. McCoy, A. J., Grosse-Kunstleve, R. W., Adams, P. D., Winn, M. D., Storoni, L. C., and Read, R. J. (2007) Phaser crystallographic software. *J. Appl. Crystallogr.* **40**, 658-674
9. Afonine, P. V., Grosse-Kunstleve, R. W., Echols, N., Headd, J. J., Moriarty, N. W., Mustyakimov, M., Terwilliger, T. C., Urzhumtsev, A., Zwart, P. H., and Adams, P. D. (2012) Towards automated crystallographic structure refinement with phenix.refine. *Acta Crystallogr. D Biol. Crystallogr.* **68**, 352-367
10. Emsley, P., Lohkamp, B., Scott, W. G., and Cowtan, K. (2010) Features and development of Coot. *Acta Crystallogr. D Biol. Crystallogr.* **66**, 486-501
11. Pettersen, E. F., Goddard, T. D., Huang, C. C., Couch, G. S., Greenblatt, D. M., Meng, E. C., and Ferrin, T. E. (2004) UCSF Chimera--a visualization system for exploratory research and analysis. *J. Comput. Chem.* **25**, 1605-1612
12. Josephs, T. M., Liptak, M. D., Hughes, G., Lo, A., Smith, R. M., Wilbanks, S. M., Bren, K. L., and Ledgerwood, E. C. (2013) Conformational change and human cytochrome c function: mutation of residue 41 modulates caspase activation and destabilizes Met-80 coordination. *J. Biol. Inorg. Chem.* **18**, 289-297

**Table S1.** Biophysical properties of cytochrome *c* variants.

| | Soret band max (nm) | Redox potential (mV ± SD, n=3-6) | Alkaline transition (p*K* ± SD, n=3) | $T_m$ (°C ± SD, n=3) |
|---|---|---|---|---|
| WT | 410 | 226 ± 3 [a] | 9.3 ± 0.4 [a] | 66.7 ± 0.2 [a] |
| G41S | 409 | 227 ± 4 [a] | 7.8 ± 0.3 [a] | 66.6 ± 0.8 [a] |
| G41A | 410 | 244 ± 3 [a] | 8.1 ± 0.5 [a] | 67.9 ± 0.5 [a] |
| G41T | 407 | 195 ± 1 [a] | 6.7 ± 0.2 [a] | 67.6 ± 0.5 [a] |
| Y48H | 408 | 221 ± 3 | 8.4 ± 0.1 | 69.5 ± 1.3 |
| A51V | 409 | 222 ± 3 | 8.4 ± 0.3 | 70.5 ± 0.9 |
| R38W | 408 | 189 ± 5 | ND | 69.7 ± 0.4 |

[a] Josephs et al 2013 (12)
$T_m$ melting temperature
ND not determined

**Table S2.** X-ray data collection, reduction and refinement statistics.

| | **Cytochrome *c* Y48H** | **Cytochrome *c* G41T** |
|---|---|---|
| **Data collection** | | |
| Beamline | Australian Synchrotron MX2 | Australian Synchrotron MX2 |
| Wavelength (Å) | 0.954 | 0.954 |
| Space Group | $P12_11$ | $P22_12_1$ |
| Unit cell a, b, c (Å)<br>α, β, γ (°) | 57, 36, 60<br>90, 116, 90 | 65, 76, 233<br>90, 90, 90 |
| [a] Resolution (Å) | 54.04 – 1.60 Å (1.62-1.60) | 77.61 – 2.70 (2.83-2.70) |
| [a] Unique reflections | 26,388 (1,364) | 32,753 (4,277) |
| [a] Redundancy | 3.7 (2.6) | 7.1 (7.3) |
| [a] Completeness (%) | 90.1 % (90.6) | 100.0 % (100.0) |
| [a,b] $R_{merge}$ | 0.095 (0.362) | 0.199 (0.713) |
| [a] I/σI | 10.4 (2.6) | 7.9 (2.6) |
| [a,c] $CC_{1/2}$ | 0.991 (0.678) | 0.990 (0.852) |
| **Refinement** | | |
| [d] $R_{work}$ / $R_{free}$ | 0.173 / 0.193 | 0.215 / 0.249 |
| Ramachandran favorable (%) | 97.1 | 96.6 |
| Ramachandran outliers (%) | 0 | 0 |
| Rotamer outliers (%) | 0 | 0 |
| Clashscore | 3.12 | 4.35 |
| R.m.s. deviation in bond lengths (Å) | 0.011 | 0.004 |
| R.m.s. deviation in bond angles (°) | 0.950 | 0.700 |
| Average B factor ($Å^2$) | 21.7 | 29.7 |
| PDB ID | 5EXQ | 6ECJ |

[a] Highest resolution shell is shown in parentheses.
[b] $R_{merge} = \Sigma_{hkl} \Sigma_i |I_i - \langle I \rangle| / -\Sigma_{hkl} \Sigma I_i$, where $I_i$ is the intensity of the $i^{th}$ observation, $\langle I \rangle$ is the mean intensity of the reflection and the summations extend over all unique reflections (hkl) and all equivalents (i), respectively.
[c] $CC_{1/2}$ is the correlation coefficient of the half datasets.
[d] $Rwork = \sum_{hkl} | |F_{obs}| - |F_{calc}| | / \sum_{hkl} |F_{obs}|$, where $F_{obs}$ and $F_{calc}$ is the observed and the calculated structure factor, respectively. $R_{free}$ is the cross-validation R factor for the test set of reflections (5 % of the total) omitted in model refinement.

**Table S3. Average RMSD ± SD (Å) for all MD simulations**

| | FeII_1 | FeII_2 | FeIII_1 | FeIII_2 |
|---|---|---|---|---|
| WT | 1.56 ± 0.14 | 1.69 ± 0.22 | 1.87 ± 0.29 | 1.63 ± 0.12 |
| G41S | 1.57 ± 0.18 | 2.25 ± 0.43 | 2.77 ± 0.47 | 1.78 ± 0.22 |
| G41A | 1.78 ± 0.22 | 1.54 ± 0.12 | 1.75 ± 0.22 | 1.63 ± 0.24 |
| G41T | 1.78 ± 0.22 | 1.74 ± 0.16 | 1.94 ± 0.19 | 1.61 ± 0.14 |
| Y48H | 1.65 ± 0.20 | 1.57 ± 0.25 | 2.18 ± 0.33 | 1.72 ± 0.19 |
| A51V | 1.98 ± 0.27 | 1.78 ± 0.30 | 1.97 ± 0.38 | 1.93 ± 0.13 |

**Table S4.** Correlation of RMSF with redox potential and alkaline transition p*K*.

| | redox potential (E°, mV)) | | | | alkaline transition (p*K*) | | | |
|---|---|---|---|---|---|---|---|---|
| residues | 7-102 | 20-35 | 40-57 | 70-85 | 7-102 | 20-35 | 40-57 | 70-85 |
| r | 0.09 | -0.19 | 0.14 | 0.21 | 0.26 | 0.23 | -0.03 | 0.67 |
| *P* | 0.85 | 0.72 | 0.79 | 0.69 | 0.62 | 0.66 | 0.96 | 0.14 |

Pearson correlation r and *P* value for average RMSF for Cα in each region from simulation data for ferric cytochromes *c*.

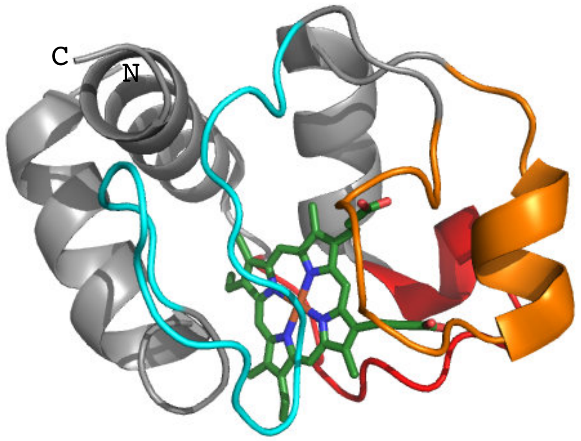


```
                   Heme
1           10   |  |  20          30          40
GDVEKGKKI FIMKCSQCHT VEKGGKHKTG PNLHGLFGRK TGQAPGYSYT
ααααααααα αααα                          βββ

50          60          70          80          90          100
AANKNKGIIW GEDTLMEYLE NPKKYIPGTK MIFVGIKKKE ERADLIAYLK KATNE
        ββ ββααααααααα                      ααα αααααααααα ααααα
```

**Figure S1.** Structure and sequence of human ferric cytochrome *c* (3ZCF). Position of the Ω loops: 19-36 in blue, 40-57 in orange, 71-85 in red. Residues involved in α helices (α) and β sheets (β) are shown.

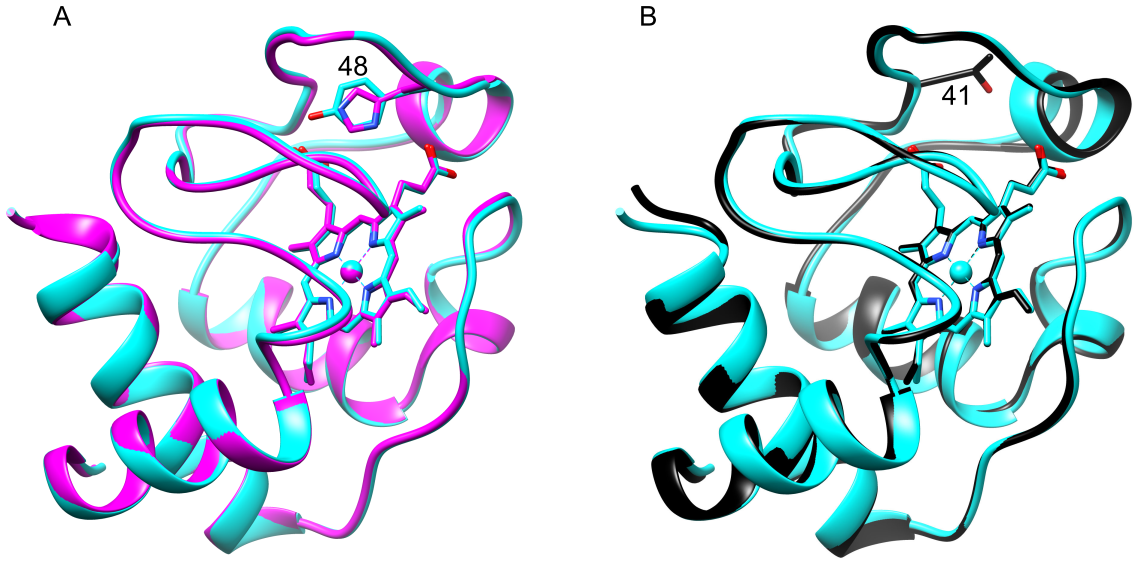


**Figure S2.** Comparison of variant human cytochrome *c* structures. (A) Alignment of Y48H (magenta, chain A, 5EXQ)) with WT (cyan, 3ZCF). (B) Alignment of G41T (Black, chain B, 6ECJ) with WT (cyan, 3ZCF). Residue 48 (A) or 41 (B) sidechain is shown.

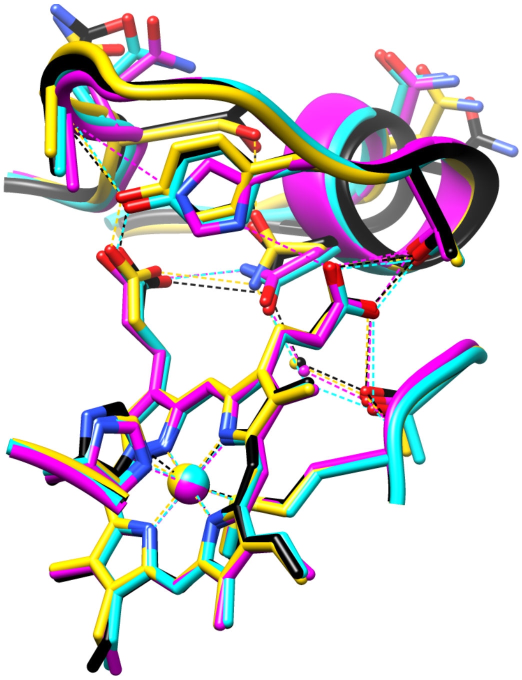

**Figure S3**. Comparison of the H-bond network of the human cytochrome *c* heme and 40-57 Ω-loop. G41T (black, chain A, 6ECJ), G41S (yellow, chain B, 3NWV (1)), WT (cyan, chain A, 3ZCF (3)), Y48H (magenta, chain A, 5EXQ). Residues 39-57 are shown.

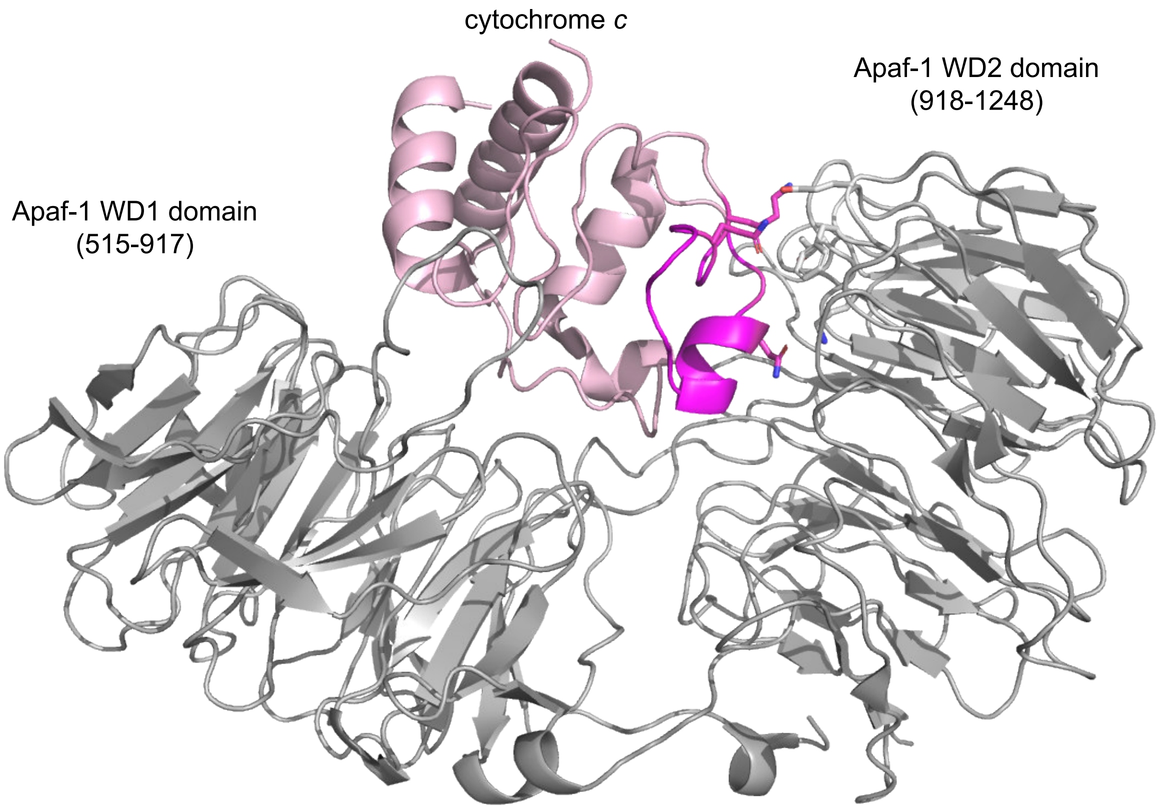


**Figure S4.** Location of cytochrome *c* between the WD1 and WD2 domains of Apaf-1. Apaf-1 in grey, equine cytochrome *c* in pale pink. The region of cytochrome *c* shown in Figure S3 is in magenta. Drawn from 5WVE.

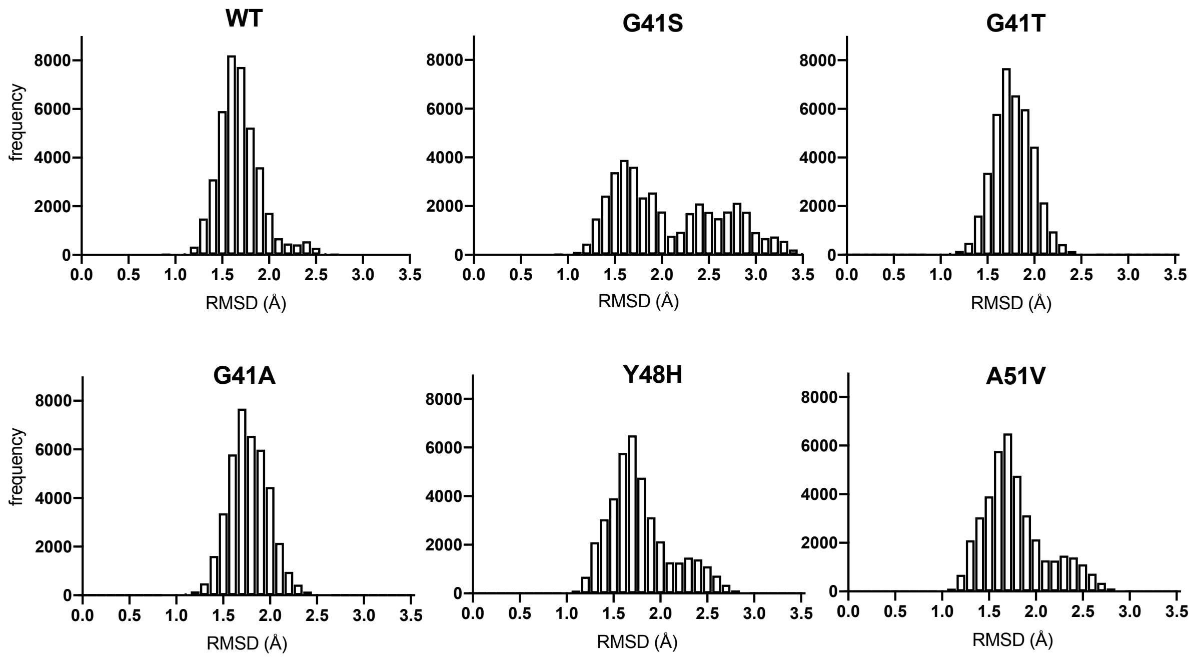


**Figure S5.** Histograms of RMSD for combined MD runs for WT cytochrome *c* and each variant.

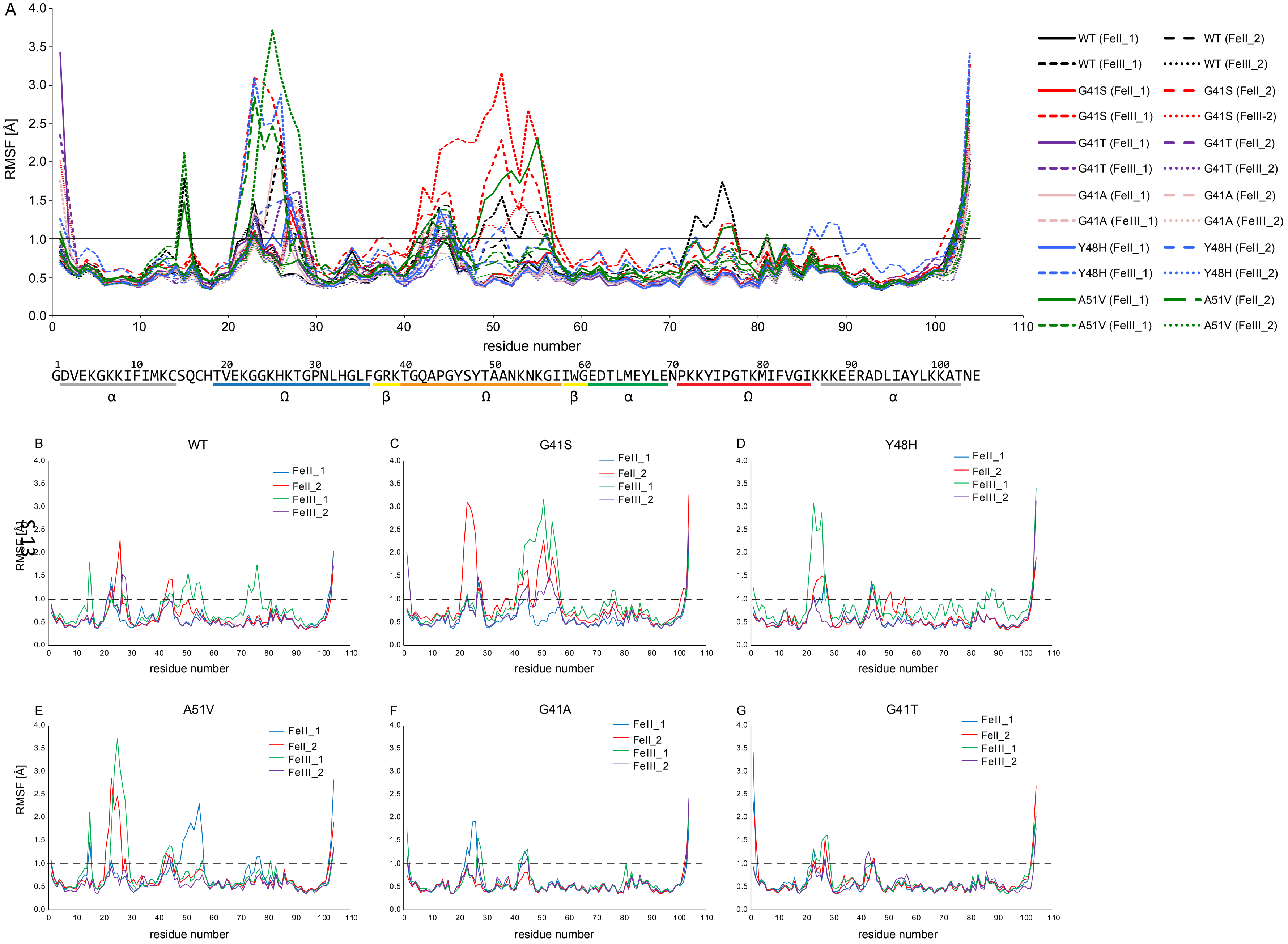




**re S6.** (A) The RMSF of Cα atoms as a function of amino acid residue for all simulations: WT cyan; Y48H magenta; G41S yellow; G41T, black; V dark green; G41A blue. The dashed line is at 1 Å. Sequence and secondary structure elements are shown below. (B-G) The RMSF of Cα atoms as ction of amino acids for each of four individual simulations run for each variant.

HEMR 105.A CGD <-> THR 49.A OG1

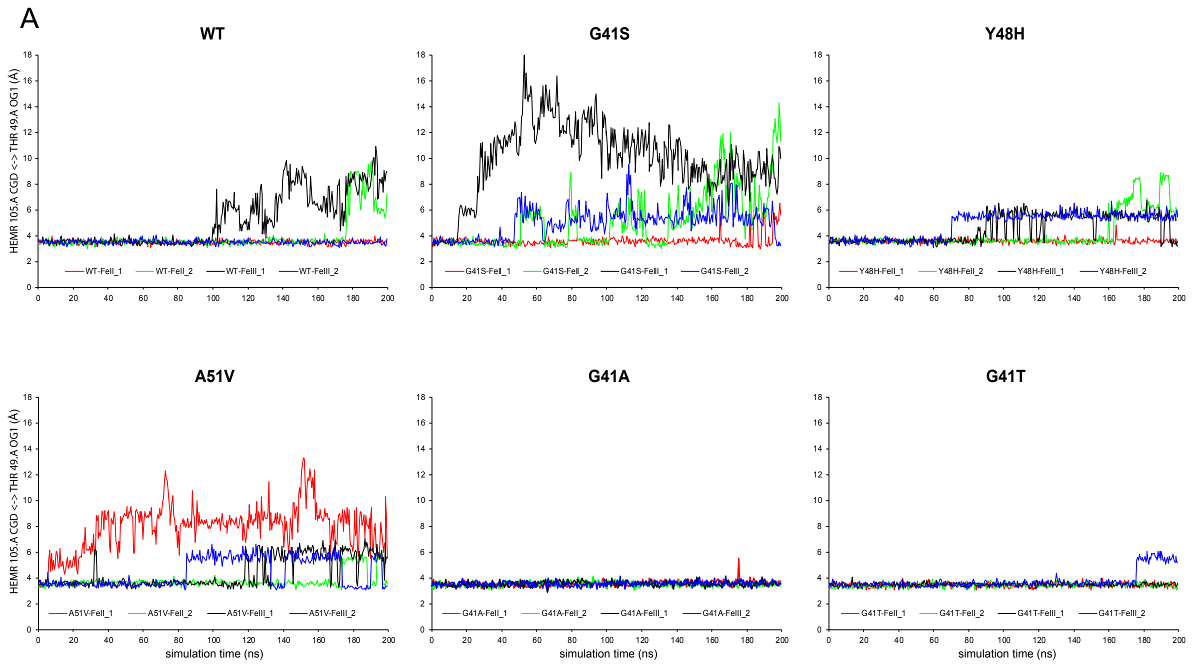


HEMR 105.A CGA <-> ASN 52.A ND2

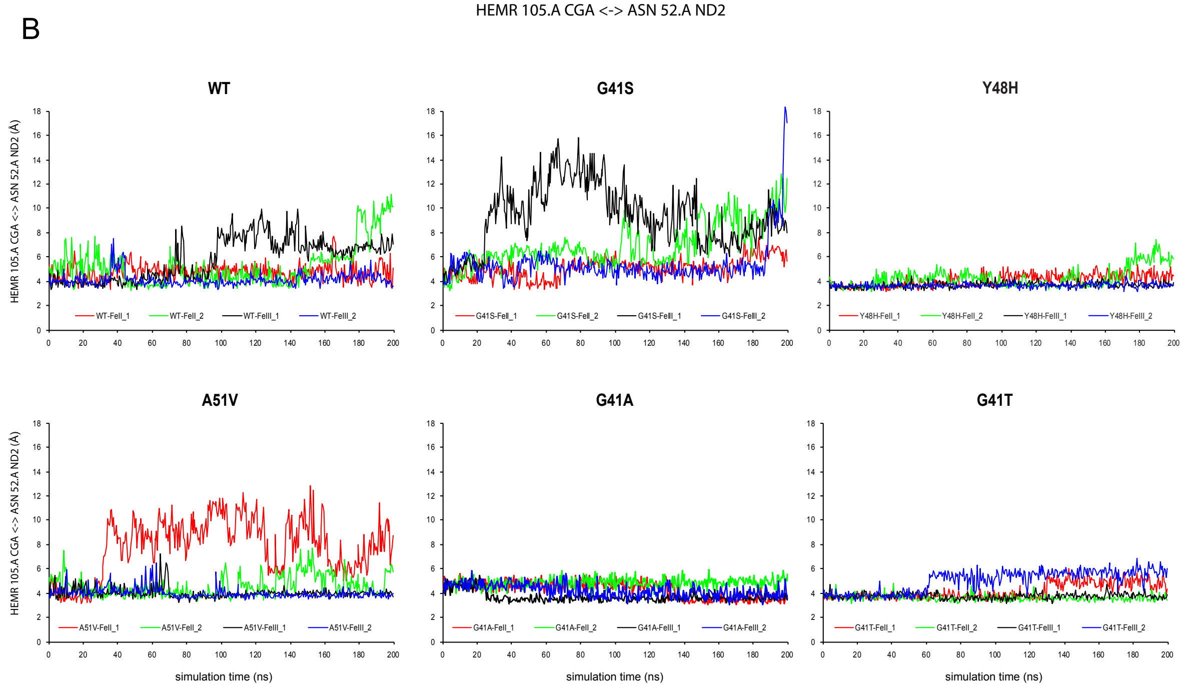

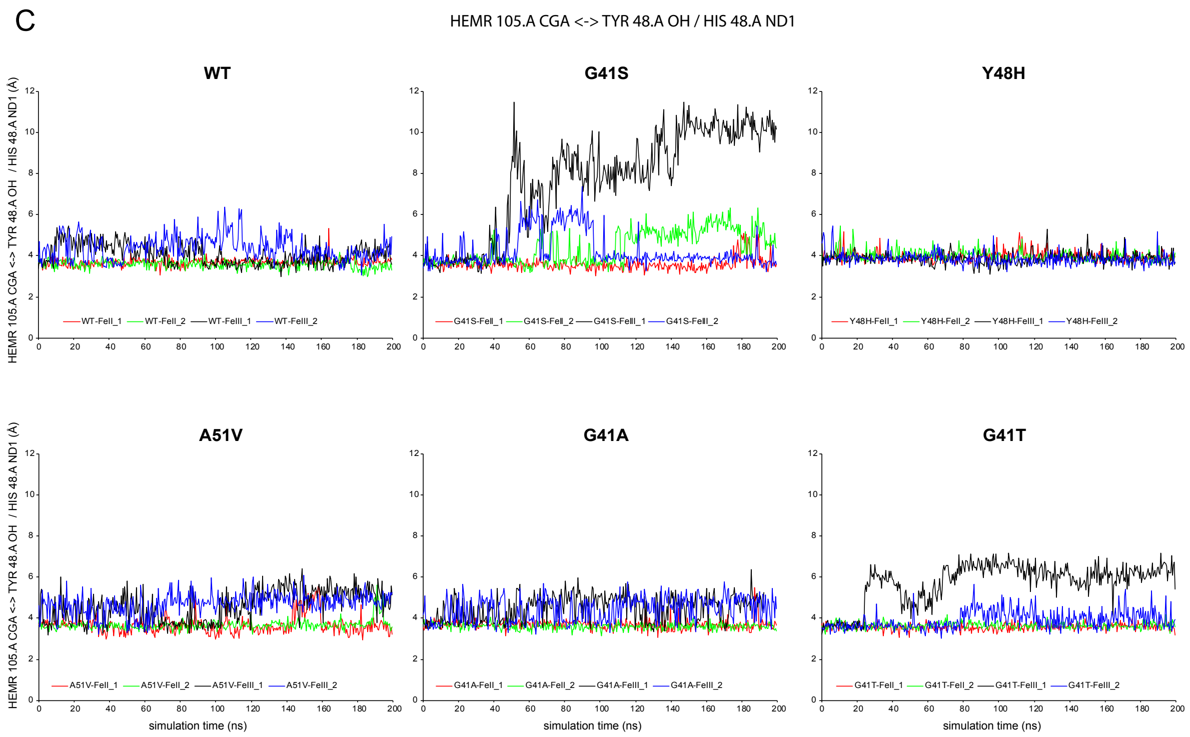


**Figure S7.** Distance from Thr49 hydroxyl to heme carboxylate CGD (A), Asn52 side chain amide nitrogen to heme propionate CGA (B), and Tyr48 hydroxyl or His48 side chain amine ND1 to heme propionate CGA (C) in simulations for WT, G41S, Y48H, A51V, G41A and G41T cytochromes *c*. Runs are colored red (FeII_1), green (FeII_2), black (FeIII_1) and blue (FeIII_2).

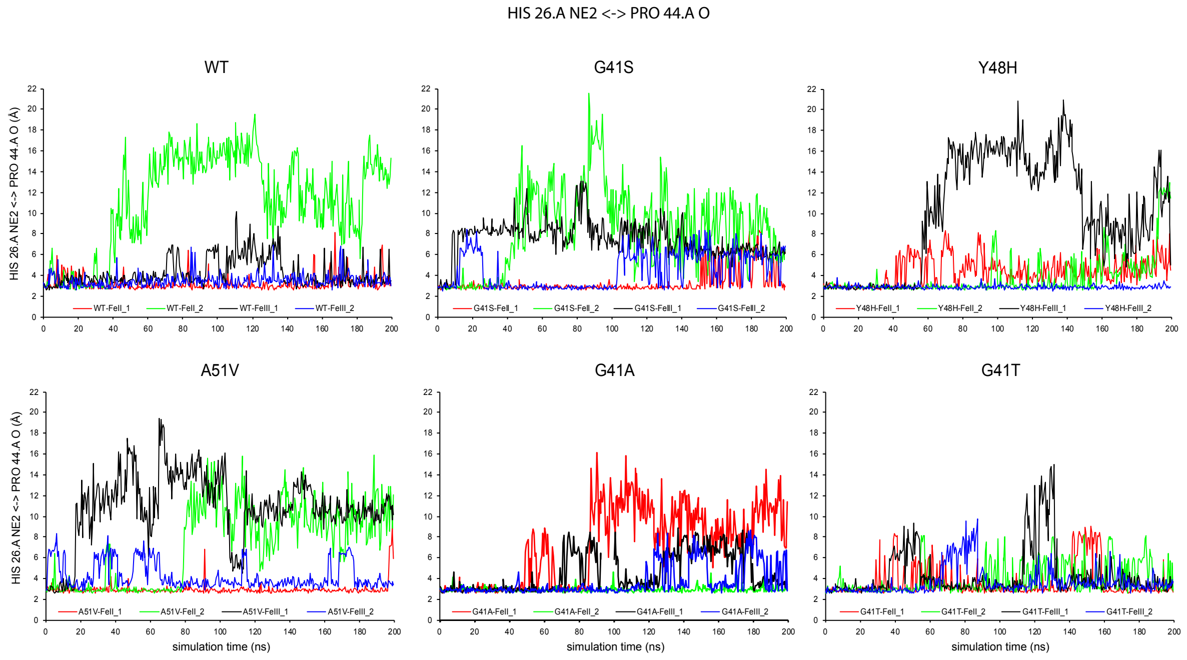


**Figure S8.** Distance from His26 amine NE2 to Pro44 carbonyl in simulations for WT, G41S, Y48H, A51V, G41A and G41T cytochromes *c*. Runs are colored red (FeII_1), green (FeII_2), black (FeIII_1) and blue (FeIII_2).

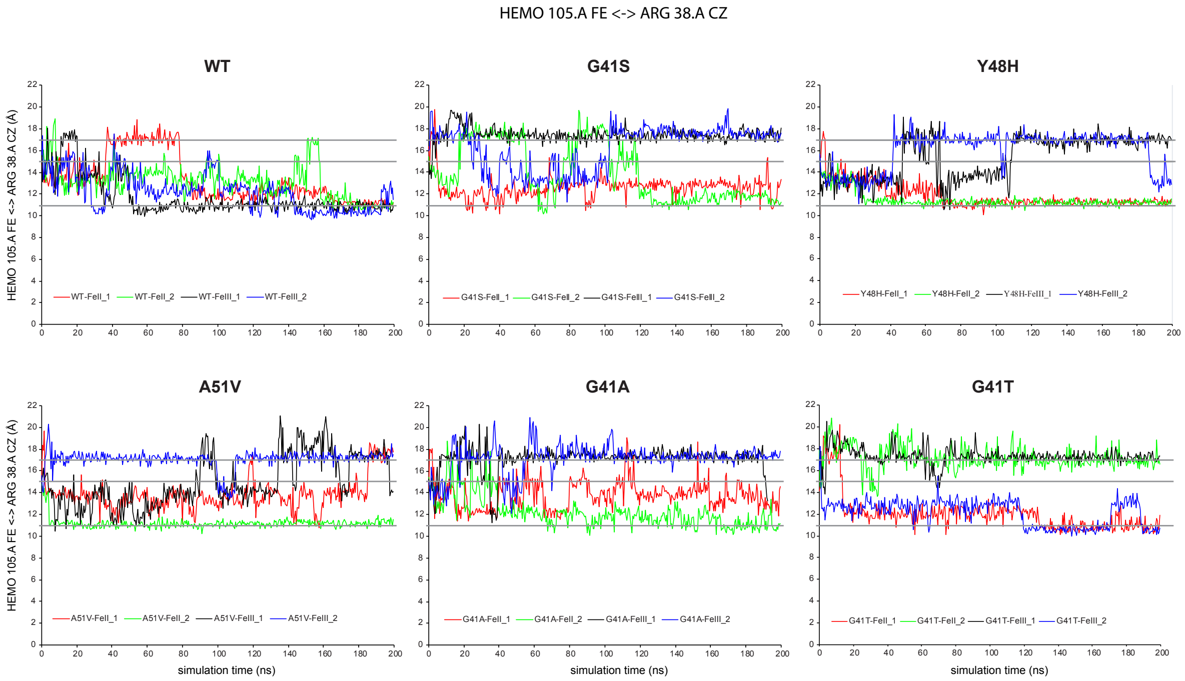


**Figure S9.** Distance from Arg38 guanidino carbon (CZ) to the heme Fe in simulations for WT, G41S, Y48H, A51V, G41A and G41T cytochromes *c*. The grey lines show the three "stable" positions with distances of 11, 15 and 17 Å. Runs are colored red (FeII_1), green (FeII_2), black (FeIII_1) and blue (FeIII_2).

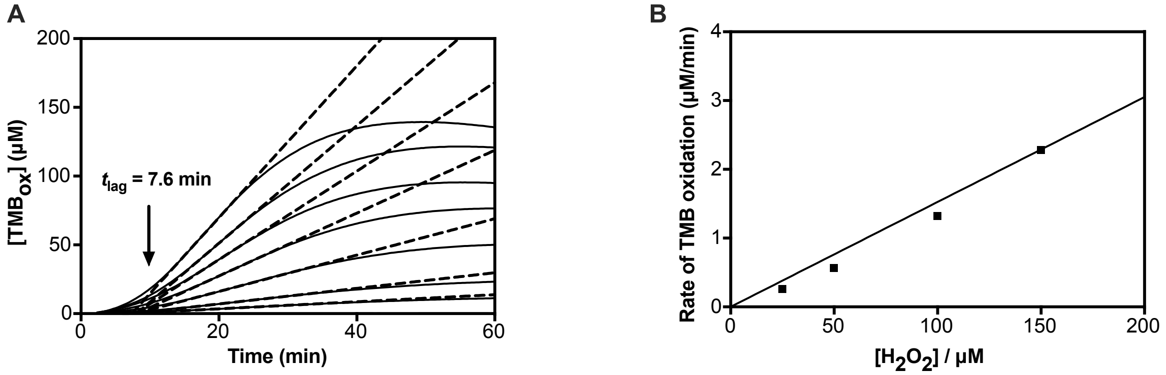


**Figure S10.** Kinetics of the peroxidase reaction for R38W cytochrome *c* with TMB at pH 5.4. (A) The plots of [$TMB_{ox}$] versus time show a lag phase followed by a steady state phase that can be fitted to a straight line (dashed). These lines ([$H_2O_2$] = 25-300 μM) intersect at a single point defining the lag time. (B) Plot of the slope versus [$H_2O_2$] is almost linear, allowing the second order rate constant to be approximated. Protein was 5 μM in heme.

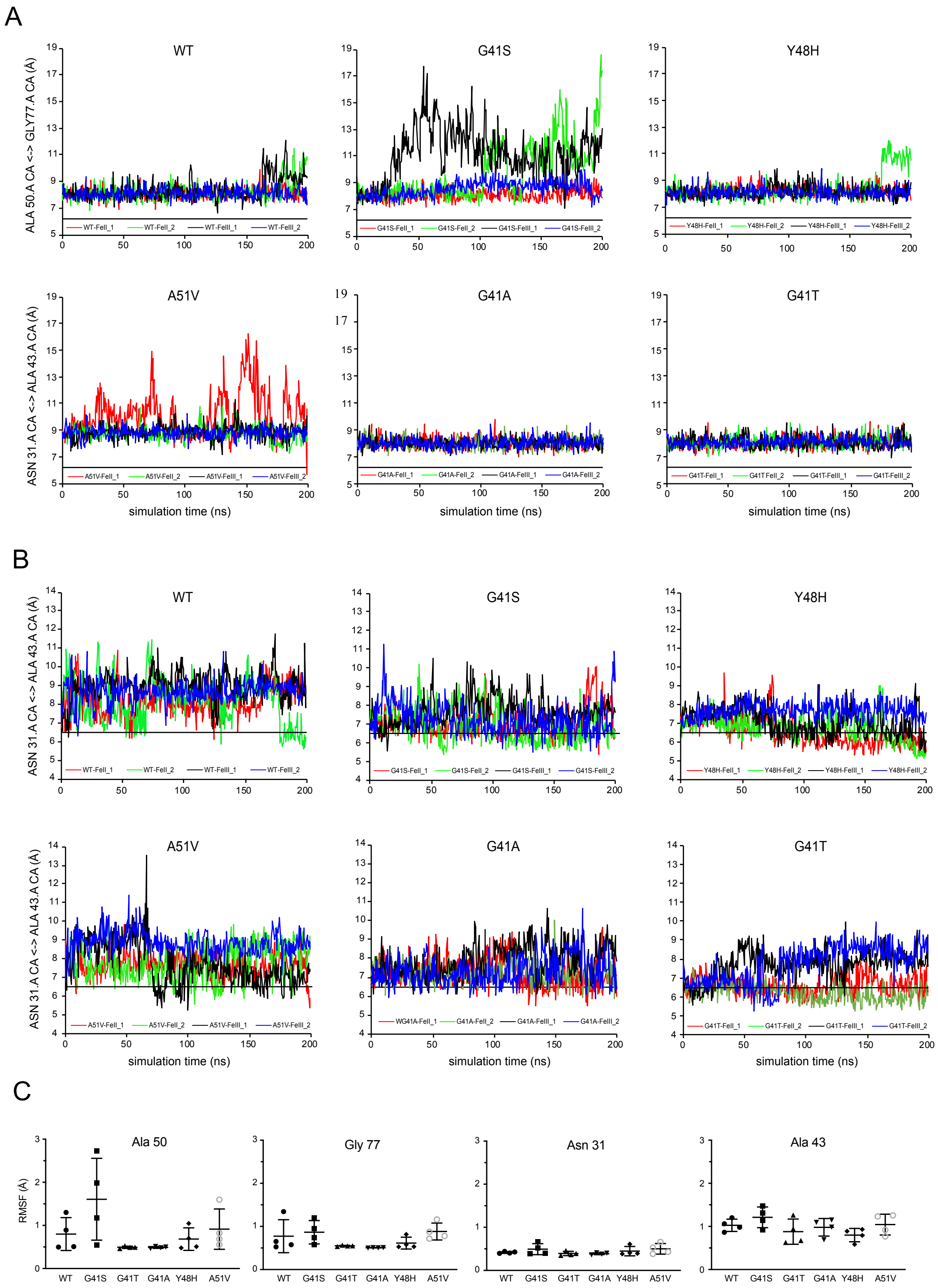


**Figure S11.** (A) Distance from Ala50 Cα to Gly77 Cα for WT, G41S, Y48H, A51V, G41A and G41T cytochromes *c*. The black line shows the 6.2 Å distance defined by Bortolotti *et al* as open for cavity A. (B) Distance from Asn31 Cα to Ala43 Cα for WT, G41S, Y48H, A51V, G41A and G41T cytochromes *c*.

The black line shows the 6.5 Å distance defined by Bortolotti *et al* as open for cavity B. Runs are colored red (FeII_1), green (FeII_2), black (FeIII_1) and blue (FeIII_2). (C) RMSF for Cα of Ala50, GLy77, Asn31 and Ala43 for all runs.

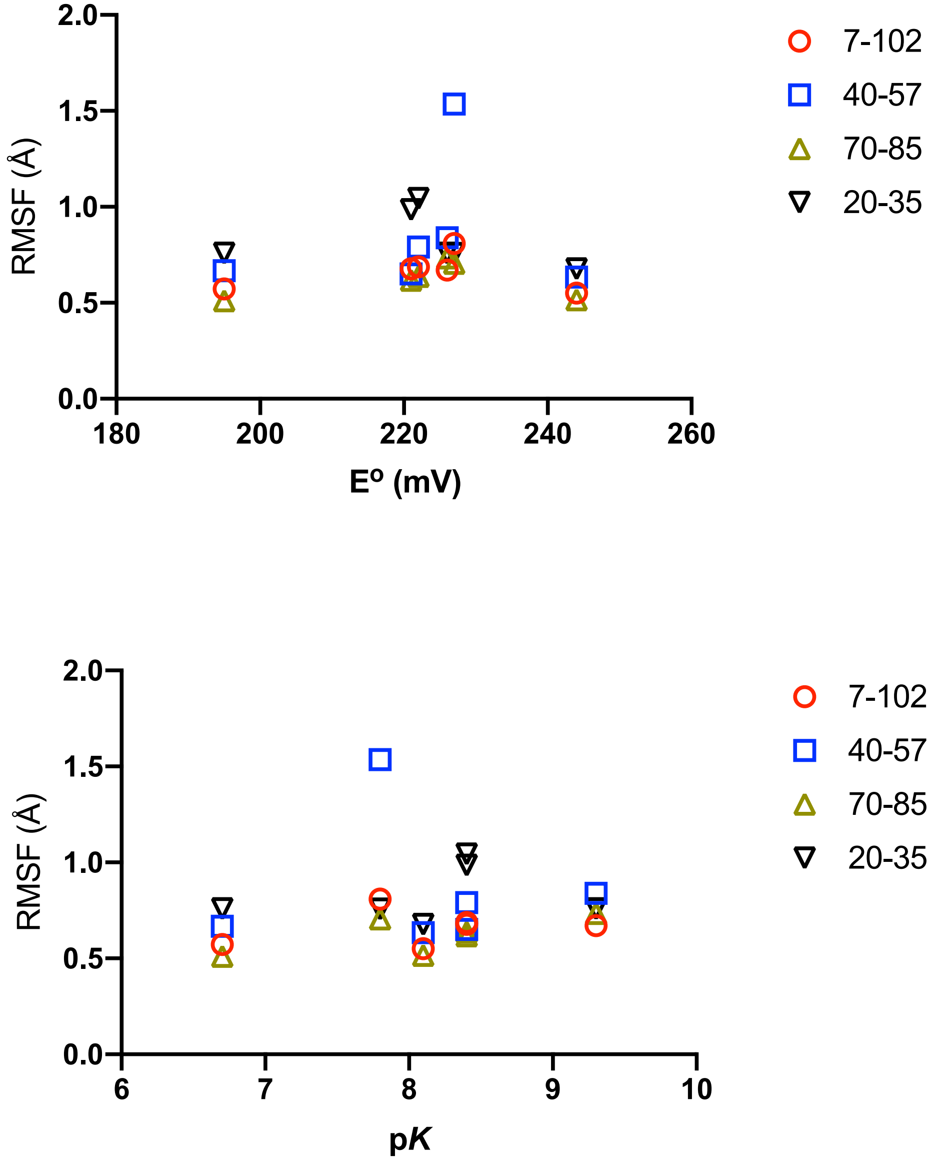


**Figure S12.** Correlation of redox potential (Eº, upper panel) or p*K* (lower panel) with RMSF for residues 7-102, the 20-35 Ω loop, the 40-57 Ω loop, and the 70-85 Ω loop.
